# The component parts of bacteriophage virions accurately defined by a machine-learning approach built on evolutionary features

**DOI:** 10.1101/2021.02.28.433281

**Authors:** Tze Y. Thung, Murray E. White, Wei Dai, Jonathan J. Wilksch, Rebecca S. Bamert, Andrea Rocker, Christopher J Stubenrauch, Daniel Williams, Cheng Huang, Ralf Schittelhelm, Jeremy J. Barr, Eleanor Jameson, Sheena McGowan, Yanju Zhang, Jiawei Wang, Rhys A. Dunstan, Trevor Lithgow

**Affiliations:** Infection & Immunity Program, Biomedicine Discovery Institute and Department of Microbiology, Monash University, Clayton, Australia; Centre to Impact AMR, Monash University, Clayton, Australia; School of Computer Science and Information Security, Guilin University of Electronic Technology, Guilin 541004, China; Monash Proteomics & Metabolomics Facility, Biomedicine Discovery Institute and Department of Biochemistry and Molecular Biology, Monash University, Clayton, Australia; School of Biological Sciences, Monash University, Clayton, Australia; School of Life Sciences, University of Warwick, Gibbet Hill Road, Coventry CV4 7AL, UK

**Keywords:** antimicrobial resistance, phage therapy, bacteriophage, artificial intelligence

## Abstract

Antimicrobial resistance (AMR) continues to evolve as a major threat to human health and new strategies are required for the treatment of AMR infections. Bacteriophages (phages) that kill bacterial pathogens are being identified for use in phage therapies, with the intention to apply these bactericidal viruses directly into the infection sites in bespoke phage cocktails. Despite the great unsampled phage diversity for this purpose, an issue hampering the roll out of phage therapy is the poor quality annotation of many of the phage genomes, particularly for those from infrequently sampled environmental sources. We developed a computational tool called STEP^3^ to use the “evolutionary features” that can be recognized in genome sequences of diverse phages. These features, when integrated into an ensemble framework, achieved a stable and robust prediction performance when benchmarked against other prediction tools using phages from diverse sources. Validation of the prediction accuracy of STEP^3^ was conducted with high-resolution mass spectrometry analysis of two novel phages, isolated from a watercourse in the Southern Hemisphere. STEP^3^ provides a robust computational approach to distinguish specific and universal features in phages to improve the quality of phage cocktails, and is available for use at http://step3.erc.monash.edu/.

**IMPORTANCE:** In response to the global problem of antimicrobial resistance there are moves to use bacteriophages (phages) as therapeutic agents. Selecting which phages will be effective therapeutics relies on interpreting features contributing to shelf-life and applicability to diagnosed infections. However, the protein components of the phage virions that dictate these properties vary so much in sequence that best estimates suggest failure to recognize up to 90% of them. We have utilised this diversity in evolutionary features as an advantage, to apply machine learning for prediction accuracy for diverse components in phage virions. We benchmark this new tool showing the accurate recognition and evaluation of phage components parts using genome sequence data of phages from under-sampled environments, where the richest diversity of phage still lies.

## INTRODUCTION

Antimicrobial resistance (AMR) has risen to prominence as a major threat to human health (1, 2) and new strategies are required for the treatment of AMR infections (3–5). For example, the Centers for Disease Control and Prevention have identified several species of microbes as “Urgent” threats to human health by virtue of their AMR phenotypes, including *Escherichia coli* and *Enterococcus faecalis*. As another prime example of one of these, the carbapenem-resistant *Enterobacteriaceae* (CRE), *Klebsiella pneumoniae* infections represent a key target for new therapeutics to treat AMR infections (3–5). Bacteriophages (phages) that kill bacterial pathogens such as *Klebsiella* are being identified for use in phage therapies, with the intention to apply these bactericidal viruses directly into the infection sites. Careful consideration is needed in selecting the phages for use in therapeutic cocktails (4–6), considerations made difficult because annotation of phage genomes is poor (7, 8), potentially obscuring phages with therapeutic potential. For example, while structural motifs are now known (9) that will promote phage virion stability (i.e. shelf-life), only with correct annotation of the major capsid, minor capsid and other proteins involved can structural motifs be identified and evaluated.

Phage therapy has re-emerged because of its potential treatment for antimicrobial-resistant infections, and a common protocol for treatments is to select two or more phages for combination into a treatment cocktail (4–6). An ongoing issue is the establishment of criteria used for selection of appropriate phages for a cocktail, to enhance production and maximize efficacy, and to circumvent issues of phage-resistance and collateral induction of further drug-resistance in the infection sites (4, 6). The phages used for phage therapy are *Caudovirales* conforming to a blue-print of an icosahedral protein capsid housing the phage genome, and a tail composed of 20-40 protein components (10). The tail of these phages can be considered as a complex piece of molecular machinery, with component parts of the tail recognizing and docking to a species-specific receptor on the host bacterium (11, 12). Penetration of the host cell envelope depends on other components of the tail, which can have enzymatic functions to locally hydrolyze each of the distinct layers of the bacterial envelope (12–14). An ultimate goal for the development of personalized phage therapy is the recognition of all of these components from genome sequence data, so that bespoke phage could be selected for specific therapeutic purposes (5, 6). However, the annotation of phage genomes is poor, potentially obscuring important features contributed by some component parts such as contributions to virion stability and shelf-life, host-range and bacterial cell lysis (7, 8, 15).

## RESULTS AND DISCUSSION

Currently phage genomes are assessed by tools such as multiPhATE (15) which provides a bioinformatics pipeline for functional annotation using sequence-based queries. The annotation accuracy of multiPhATE is limited by the extreme sequence diversity in phage genomes, likely due to the rapid evolutionary rates of phages (16). This limitation has been addressed to some extent with a neural network-based predictor iVIREONS (17) and further tools such as PVPred (18), PVP-SVM (19), PhagePred (20), Pred-BVP-Unb (21) and PVPred-SCM (22). However, recent evaluation of these tools in phage protein prediction showed less than satisfactory performance (23). We developed an ensemble predictor, STEP^3^, to accurately call the protein components of phage virions and visualize their predicted function-based relationships (Fig. 1).

**Figure 1.**
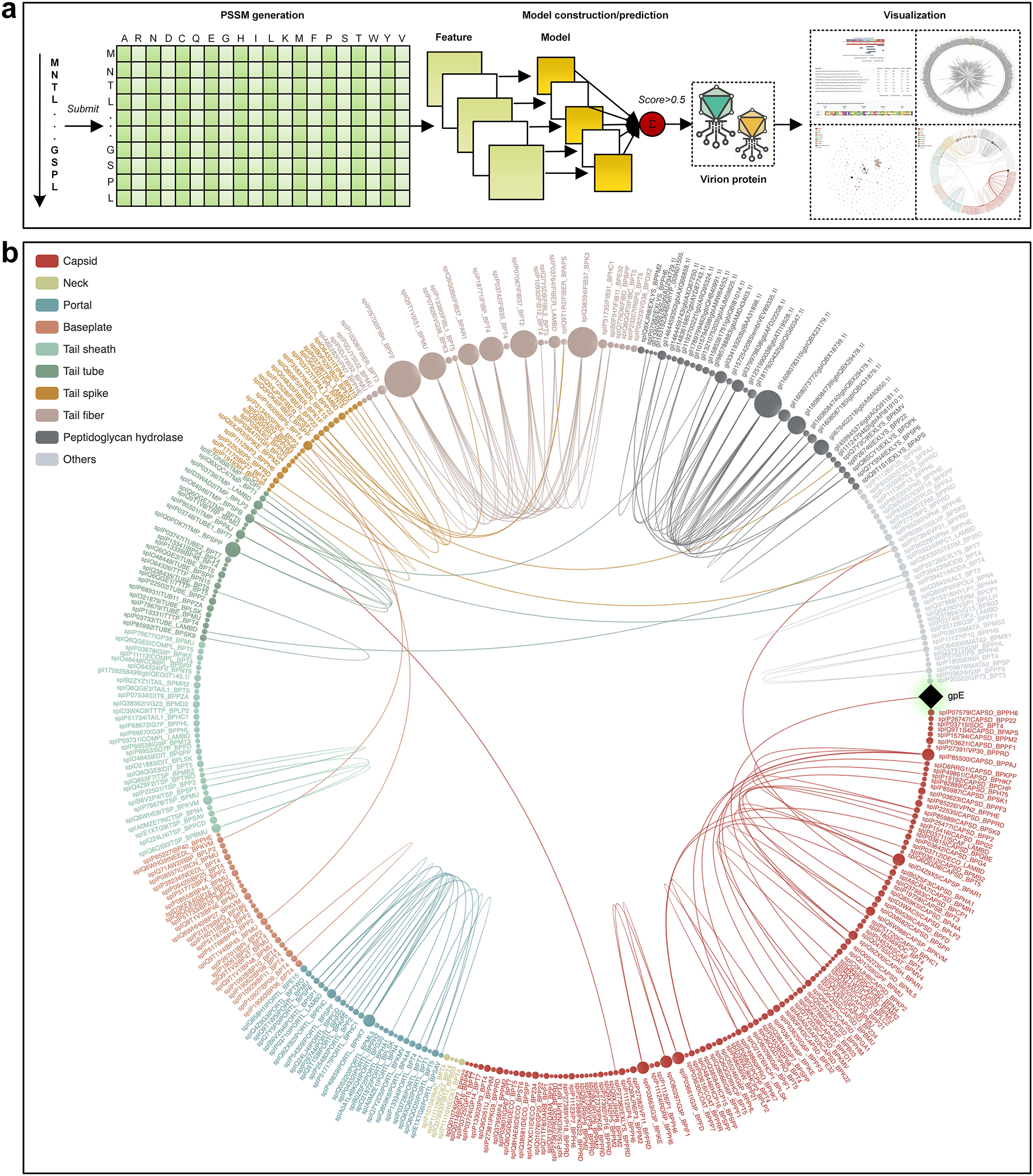
Construction and workflow for STEP^3^. (a) Graphic summarizing the construction and prediction process of STEP^3^. A set of experimentally validated virion proteins and non-virion proteins was compiled and sequence data fed into five PSSM models, including AAC-PSSM(60), PSSM-composition(61), DPC-PSSM(60), AADP-PSSM(60), and a MEDP(62) model. The five individual models were trained based on five balanced subsets, and their prediction scores were averaged to obtain an ensemble model. Finally, five baseline models (corresponding to five evolutionary features) were further integrated as the final ensemble model of STEP^3^ through averaging their prediction scores. Support vector machine (SVM) with a radial basis function kernel was used to train each model. This ultimately provides a prediction of a “virion protein” which would be a structural component of the phage virion. (b) STEP3 data visualization provides a means to document relationships between a protein of interest. The example given is the protein component gpE from phage λ, which shows clear similarity to major capsid proteins from other phages. Structural studies confirm that despite limited sequence similarity, gpE is part of a family of major capsid proteins(9). Alternative visualization features are available in STEP^3^ (Supplementary Fig. 1).

STEP^3^ extracted information from Position-Specific Scoring Matrix (PSSM) data (Fig. 1a), an approach that tracks protein evolutionary histories (24, 25). In machine-learning evaluation of protein sequences, “evolutionary features” refer to information within the amino acid sequences that conceptually traces the evolutionary history of proteins, and their use often identifies highly informative patterns (24, 25). Indirectly, these evolutionary features effectively capture structural as well as physicochemical properties. STEP^3^ includes data visualization capabilities to document relationships between virion components where the sequence similarity is sufficiently strong to identify high confidence homologs from other phages (Fig. 1b, Supplementary Fig. 1).

There is power in integrating individual models within an ensemble framework for more robust and stable predictions: trained with an individual model alone (AAC-PSSM), predictions perform well with the 5-fold cross-validation test (Supplementary Table 1), but ranked only fourth using the independent test (Supplementary Table 2). In contrast, combined with other models into the ensemble model of STEP^3^, to draw on the best elements from all of the individual models (Fig. 1a), the best prediction performance ranking was achieved (Supplementary Tables 1, 2). In benchmarking against other available predictors, the ensemble STEP^3^ achieved an improved performance, with the highest sensitivity (SN = 0.896), accuracy (ACC = 0.891), F-value (0.891) and Matthews Correlation Coefficient (MCC = 0.781) using the independent test (Supplementary Table 3). The superior performance of STEP^3^ can be attributed to the integration of more informative evolutionary features, as well as the comprehensive and up-to-date training dataset using experimentally verified inputs. It is worth noting that the BLAST-based predictor, which represents the mode used for genome annotation had the lowest accuracy (ACC) and F-value. This prediction bias is reflected by the extremely unbalanced sensitivity (the lowest) and specificity (the highest) scores, so that the BLAST-based predictor tended to predict positive samples as being negative. This quantifies and evidences past observations that pairwise sequence matching methods struggle to predict phage proteins (25).

As initial case studies we drew on three accounts published after STEP^3^ was trained, where phages had been discovered, the genome sequence data deposited for public access, and the protein composition virions had been determined by mass spectrometry. The mass spectrometry data is crucial as it enables a discrimination between false positive (FP; predicted but not present by mass spectrometry of the virion) and true positive (TP; predicted and found present by mass spectrometry of the virion). Phage vB_EfaS_271 infects *Enterococcus faecalis* (26), phage vB_PatM_CB7 infects *Pectobacterium atrosepticum* (27), and phage vB_Eco4M-7 infects enteropathogenic *Escherichia coli* (28). STEP^3^ was benchmarked against equivalent predictors: PVPred, PVP-SVM, PVPred-SCM and Pred-BVP-Unb (Fig. 2). STEP^3^ provided the greatest set of true positive predictions for each of the three phages, predicting 9 of the 12 virion components for phage vB_EfaS_271, 23 of the 26 protein components for phage vB_PatM_CB7 and 24 out of 33 components of the phage vB_Eco4M-7 virions. Making low FP predictions on each phage, STEP^3^ maintained a good balance between TP and FP and showcased robust prediction performance across the test cases. In the case of phage vB_PatM_CB7, where mass spectrometry data had shown the number of non-virion proteins is more than eight times as many as that of virion proteins, STEP^3^ generated an equal number of FP as that of TP. In this extreme case, STEP^3^ correctly predicts 23 out of 26 virion proteins with a false positive rate of 10.1% (23/227).

**Figure 2.**
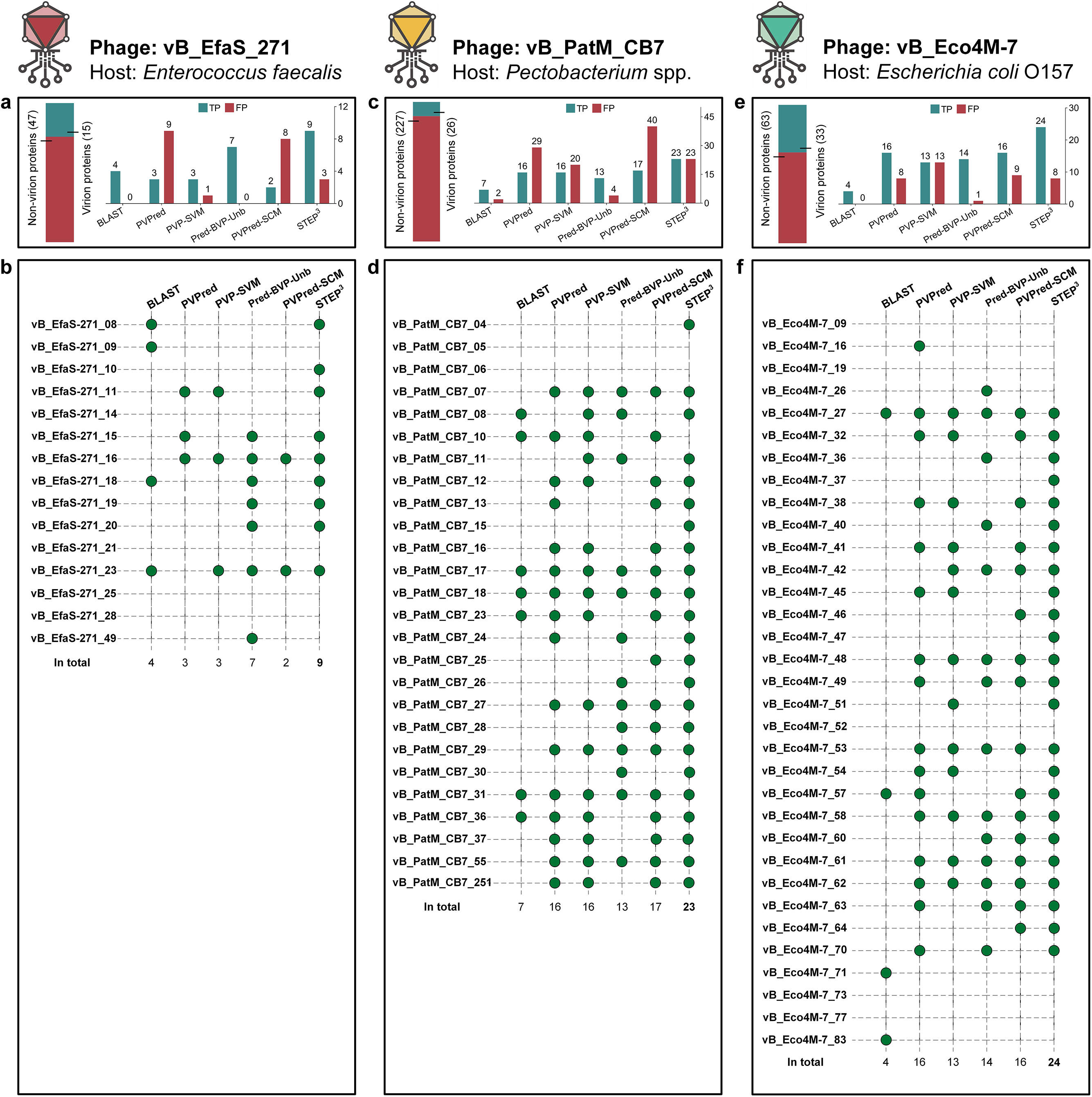
Prediction details from STEP^3^ and other tools. (a) For phage vB_EfaS_271, horizontal bars denote the number of virion and non-virion proteins. The bar chart counts the virion proteins correctly retrieved as true positives (TP), i.e. confirmed by mass spectrometry (26), and non-virion proteins mistakenly predicted as virion proteins (denoted by false positives, FP). (b) For each protein in the phage vB_EfaS_271 virion defined by mass spectrometry, a green circle represents a successful hit by a predictor. (c) For phage vB_PatM_CB7, the bar chart counts the virion proteins correctly retrieved as TP and non-virion proteins mistakenly predicted as FP. (d) Detailed predictions from STEP^3^ and other tools for vB_PatM_CB7 virion proteins defined by mass spectrometry (27). (e) For phage vB_Eco4M-7, the bar chart counts the virion proteins correctly retrieved as TP and non-virion proteins mistakenly predicted as FP. (f) Detailed predictions from STEP^3^ and other tools for vB_PatM_CB7 virion proteins defined by mass spectrometry (28).

Oftentimes candidate phages that kill pathogens are isolated from hospital waste-water sources for their use in phage therapy (29, 30). This raises the issue of potential over-sampling of a common environmental source (*i.e*. wastewater) for phages, potentially limiting discovery of other, valuable phages and also potentially biasing the capability of predictors like STEP^3^. Therefore, as a further proof of principle test for STEP^3^ we sampled a natural watercourse with a strain of drug-resistant and hypervirulent *Klebsiella pneumoniae* as host. The Merri Creek, which forms a part of the larger Merri catchment, lies within Wurundjeri Woi Wurrung people’s traditional homelands. Phages isolated from two separate sampling sites were characterized initially by genome sequencing and named in Woi wurrung language Merri-merri-uth nyilam marra-natj (MMNM) and Merri-merri baany-a bundha-natj (MMBB). These names translate as “Dangerous Merri lurker” and “Merri water biter”, respectively, in English.

Comparative genomic analysis revealed *Klebsiella* phages MMNM (Supplementary Fig. 2) and MMBB (Supplementary Fig. 3) to be distinct from previously sampled phages. In the case of MMNM, some similarities can be seen to phages belonging to the *Jedunavirus* genus according to the most recent International Committee on Taxonomy of Viruses (ICTV) classification, but the branch lengths on the tree designate diversity within this small group, comprising only eight phages in the NCBI database (Fig 3a). Relatives of MMNM, isolated from hospital wastewater in Russia, showed considerable diversity in gene content and arrangement (Fig 3b). Most notably, MMNM encodes several genes that are absent in many of the other sequenced *Jedunaviruses*, including previously uncharacterised proteins MMNM_5, MMNM_6 MMNM_45, MMNM_51, MMNM_56, MMNM_57 and the putative polynucleotide kinase protein MMNM_50. Conversely, MMNM lacks the putative NHN endonuclease-like protein encoded by both vB_KpnM_FZ14 and vB_KpnM_KpV52. Sequence annotations (Supplementary Table 4) suggest that MMNM has a tail structure characteristic of *Myoviridae*, including a baseplate protein (MMNM_21), a baseplate J-like protein (MMNM_23) and the base-plate wedge protein (MMNM_26). In high resolution structural analyses of the *Myoviridae* phage T4, each virion has 6 molecules of each of these proteins and 1-3 molecules per virion of the hub proteins to which the baseplate is attached (31, 32).

**Figure 3.**
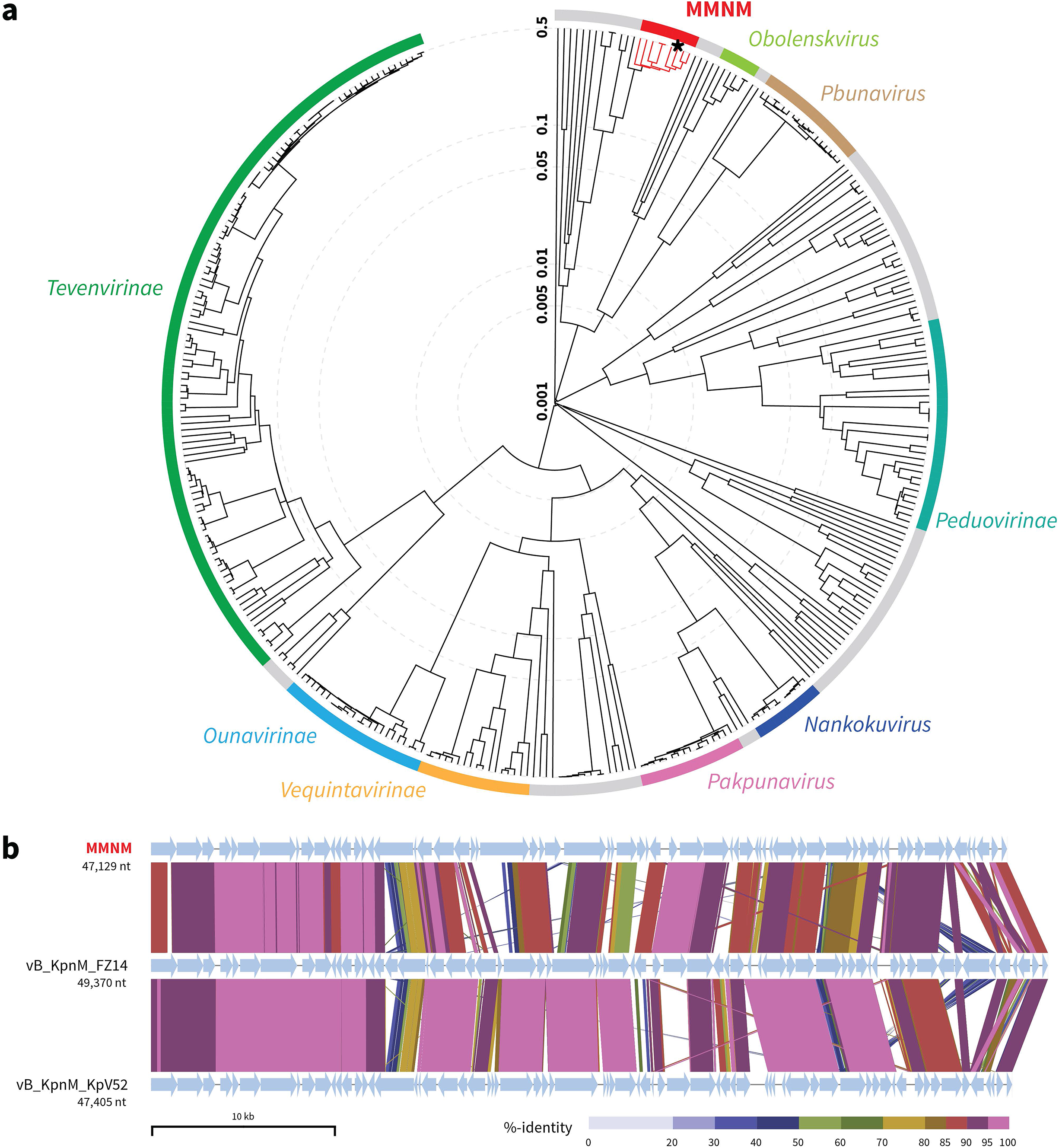
Comparative genome analysis of *Klebsiella* phage MMNM. (a) Proteomic tree analysis of *Myoviridae* that infect *Gammaproteobacteria*. The branch lengths represent genomic similarity based on normalised pairwise sequence similarity scores plotted on a logarithmic scale. The tree was constructed using sequences from the default ViPTree dataset and phage genomes listed in Table S13. Viral subfamilies or genera are highlighted in the coloured bars. Gray bars represent phages that are currently unclassified. All known members of the *Jedunavirus*, including *Klebsiella* phage MMNM (*), are highlighted in red. (b) Whole genome alignment of *Klebsiella* phage MMNM, vB_KpnM_FZ14 and vB_KpnM_KpV52. Each genome has been oriented to start with the gene encoding the putative tape measure protein. The sequences are linked by colored bars highlighting sequence identity values as shown in the key.

MMBB belongs to the *Webervirus* genus, a group of phages that exclusively target *Klebsiella* species (Supplementary Fig 3). MMBB is distinct from the other phages in this genus, with its closest relationship being to a phage isolated in China called vB_KpnS_GH-K3 (also called phage GH-K3) (33). Highlighting their differences, MMBB and GH-K3 show regions of diversity in gene content and arrangement, this is observed for the gene encoding MMBB_16, a putative AP2/HNH endonuclease previoulsy found only in a small number of other *Siphoviridae* phages including the *Escherichia* phage vB_EcoS_ESCO41 and *Escherichia* phage CJ19 (Supplementary Fig 2). Additional differences are seen in a contiguous cluster of four genes encoding hypothetical proteins (MMBB_45-MMBB_48) that are absent in GH_K3.

Phenotypic characterization of the phages on lawns of *K. pneumoniae* (Methods) showed that the plaque size for MMNM was smaller than MMBB (Fig 4a), and with liquid cultures of *K. pneumoniae* (Methods) that MMNM had a shorter latent period (L) before host cell death as determined by one-step growth curves (Fig 4b). Electron microscopy revealed that MMNM has an icosahedral head and a tail tube of ~54 nm capped with a ~30 nm baseplate to generate thick and straight tails (Fig 4c). The baseplate structure evident in MMNM (Fig 4c) is similar to that seen for the T4 phage (31), which serves as a paradigm for the *Myoviridae* (34) (Fig 4d). By contrast, MMBB has ~200nm long, slender and flexible tails (Fig 4c). The flexible, non-contractile tail tube designate MMBB as a phage of *Siphoviridae*-like viruses (Fig 4d), consistent with genome annotation data (Supplementary Table 5).

**Figure 4.**
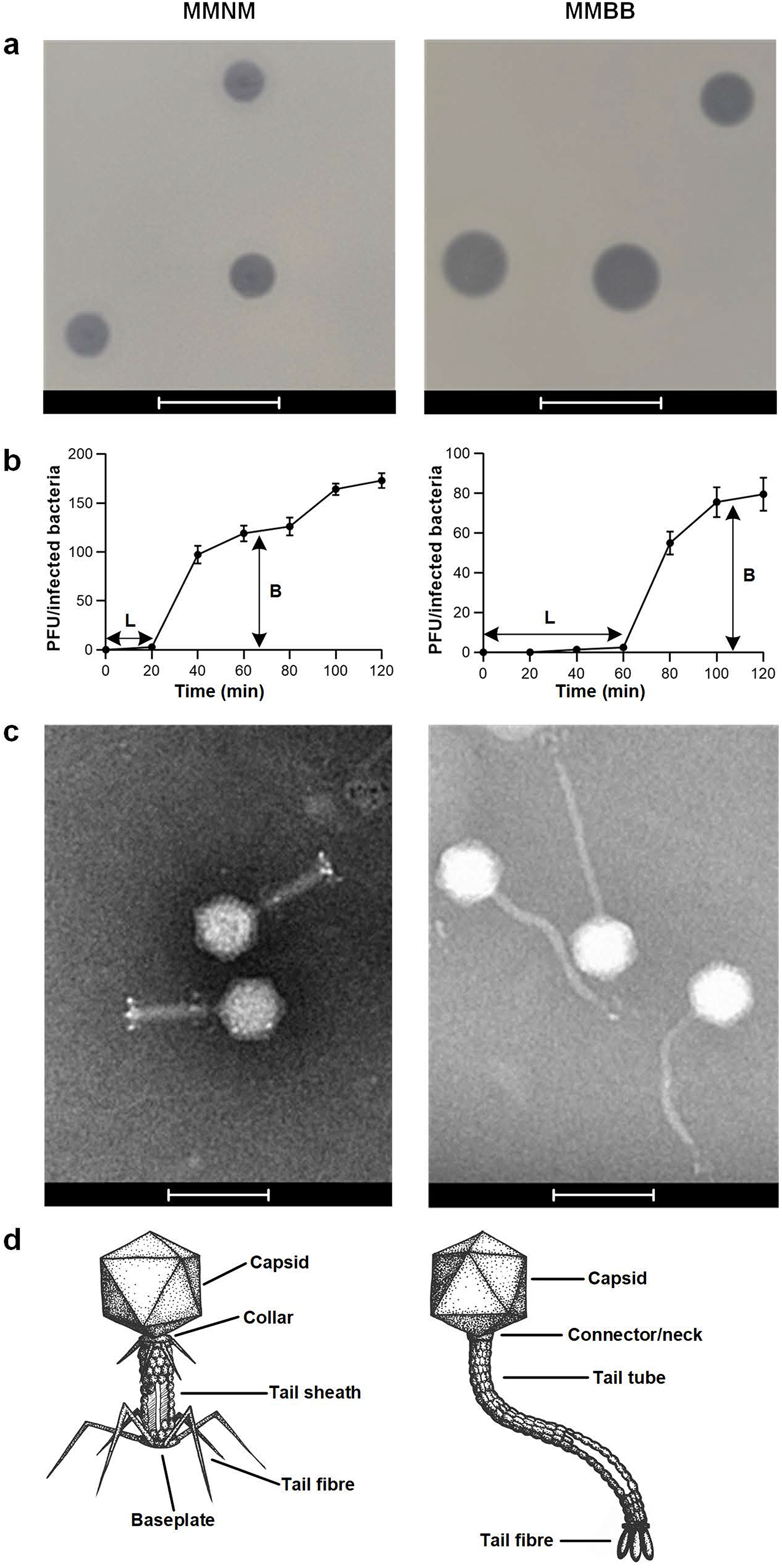
Morphological characterization of phage MMNM and MMBB. (a) Plaque morphology analysis was performed using the double overlay method. Phages MMNM and MMBB were serially diluted with SM buffer and spotted onto LB agar plates containing a top layer of soft agar and *K. pneumoniae* B5055 Δ*ompK36*. Plaque morphologies of MMNM and MMBB were determined after overnight incubation at 37°C. Scale bars represent 10 mm. (b) One-step growth curve of MMNM (left) and MMBB (right) was performed by co-incubation with the host strain for 10 min at 37°C for phage adsorption, after which the mixture was subjected to centrifugation to remove free phage particles. The resuspended cell-phage pellets were incubated at 37°C and sampled at 10 min intervals for 120 min. L, latent period; B, burst size. Data points are the mean of n=3 biologically independent samples and the error bars are the standard deviation. (c) Transmission electron micrographs of MMNM (left) and MMBB (right). The scale bars represent 100 nm. (d) Based on EM micrographs, illustrations of MMNM (left) and MMBB (right) note the cognate features in *Myoviridae* and *Syphoviridae* with annotation.

To directly test STEP^3^ prediction capability on the novel phages MMNM and MMBB, the protein components contributing structurally to the virions were determined by high-performance mass spectrometry (35, 36). To this end, samples of each virion were purified using caesium chloride gradients. The MMNM virion is composed of 25 protein components (Supplementary Table 6). Assuming a similar stoichiometry between MMNM virions and the paradigm for *Myoviridae*, phage T4 virions, the identification of the lytic transglycosylase MMNM_19 suggests that the proteomic analysis is sensitive enough to detect 3 or fewer molecules per virion (31). From evaluation of the predicted proteins within the phage genomes, together with this mass spectrometry data, the MMNM genome encodes 25 structural proteins that serve as components of the virion and 42 proteins that would be expressed after infection of the host, to drive phage replication (Fig 5a).

**Figure 5.**
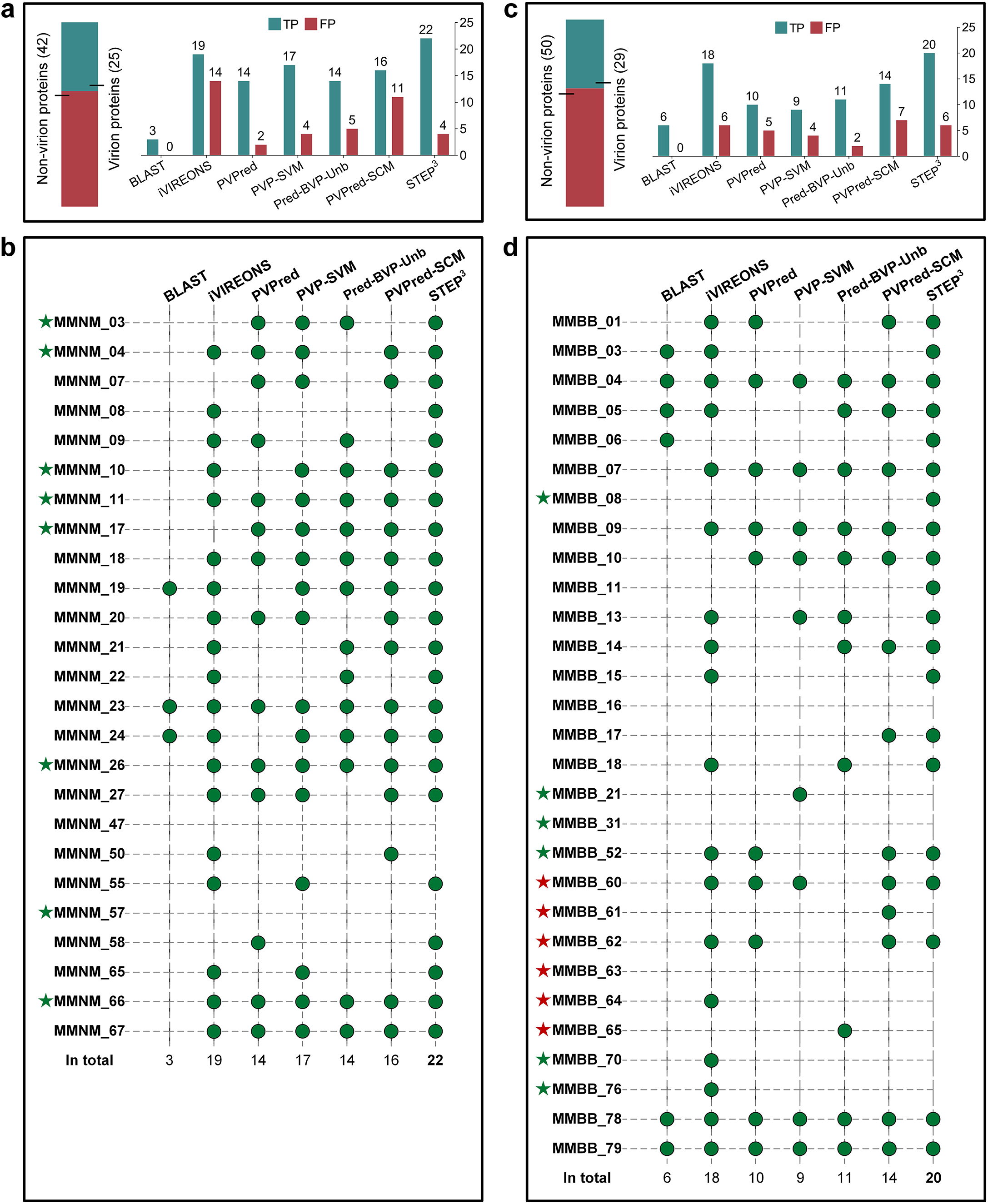
Prediction details from STEP^3^ and other tools applied to MMNM and MMBB. (a) The statistics of the prediction results on MMNM. Horizontal bars on top describe the number of virion and non-virion proteins in the phage isolates. The bar chart counts the virion proteins correctly retrieved (denoted by true positives [TP], i.e. confirmed by mass spectrometry) and non-virion proteins mistakenly predicted as virion proteins (denoted by false positives [FP]). (b) Detailed predictions from STEP^3^ and other tools for MMNM the virion proteins defined by mass spectrometry. The green circles represent a successful hit by a predictor. The green stars denote the proteins that have not previously been identified in phages. The red stars denote those with activities that have been previously identified in phages, but not previously found as protein components of purified virions. (c) Prediction statistics for MMBB. (d) Detailed predictions from STEP^3^ and other tools for MMBB virion proteins defined by mass spectrometry.

STEP^3^ successfully predicted 22 out of the 25 MMNM virion proteins (Fig 5b, Supplementary Table 7). The other predictors gave poorer outcomes with these diverse protein sequences. For example, second to STEP^3^ was iVIREONS which identified 19 virion proteins, but iVIREONS also generated the largest number of false positives, 14, consistent with its high false positive prediction rate in the independent tests (Supplementary Table 3). In one case, the initial STEP^3^ analysis made a false-negative prediction that was highly informative. The phage polynucleotide kinase (PNK) is an enzyme that has been previously assumed to be a non-virion protein, and the sequence was therefore included in that (non-virion) dataset from which STEP^3^ was trained. However, mass spectrometry identified the putative PNK protein MMNM_50 as a component of the virion Supplementary Table 6). Note, an equivalent result was achieved with the prediction for MMBB: protein MMBB_64 was detected by mass spectrometry (Supplementary Table 9) and selected by STEP^3^ (Supplementary Table 8). We suggest that for some phages the PNK remains associated with the packaged genome and is thereby incorporated within the capsid. This suggestion explains the proteomics data herein, reconciles the false-negative prediction by STEP^3^, and is consistent with the recent observation that the “gp44 ejection protein” is a virion-protein in a *Staphylococcus* phage 80α bound to genome ends and functioning as a putative PNK would to protect the DNA from degradation upon phage entry into its host (37).

High-resolution mass spectrometry of the MMBB virions showed them to be composed of 29 protein components (Supplementary Table 9). Thus, the MMBB genome encodes 29 proteins contributing structurally to the virions, and 50 non-virion proteins expressed only after infection in the host bacterium (Fig 4c). For MMBB, STEP^3^ and iVIREONS retrieved 20 and 18 virion proteins, respectively (Fig 4d, Supplementary Table 8). The other predictors achieved unsatisfactory prediction results, retrieving less than half of the 29 virion proteins.

The evolutionary features drawn on by STEP^3^ and iVIREONS are structure-informed, in that the patterns that they recognize are reflections of secondary and tertiary structure, and these patterns can also be used to suggest protein function. For example, a characteristic of the *Webervirus* has been suggested to be the presence of tail-spike proteins with polysaccharide degrading activity (38), and the sequence of MMBB_78 is suggestive of such a protein, as summarized in Supplementary Fig. 3. Conversely, pairwise sequence assessment is a poor means for recognition and characterization of virion proteins. For both MMNM and MMBB, sequence conservation alone proved the least satisfactory method for predicting phage virion proteins: the BLAST-based predictor recognized only 3 and 6 virion proteins, respectively (Fig 5b, 5d, Supplementary Tables 7, 8). This confirmed the independent test results that the BLAST-based methods commonly used for annotations are a poor means of recognizing and classifying sequence-diverse phage proteins.

Some estimates put the number of phage virions in the world at 10^31^, suggesting that there is a huge pool of phages that we know little about (39). This encourages a move towards informed bioprospecting for potentially useful phages from under-sampled environments. The effective use of these for therapy and other applications depends on a number of factors, not least of which is the sequence-based choices that must be made to identify novel phages warranting further characterization and potential development into phage therapy. We suggest that application of STEP^3^ will assist in distinguishing the specific and universal features in phages isolated from under-represented (under-sampled) geographical locations, with impact on the quality of future phage cocktails. Particularly in phage that might be highly divergent in their sequence characteristics, such as the MMNM and MMBB case studies here, STEP^3^ can predict the component parts of the virions with a confidence level well above other computational tools. The STEP^3^ toolbox is available at http://step3.erc.monash.edu/.

## METHODS

### Construction of the Klebsiella host strain

B5055 is a multidrug-resistant *K. pneumoniae* (40, 41) strain with a K2-type capsule considered indicative of hypervirulent *K. pneumoniae* (hvKp) (42). To avoid isolating phages that use the major porin for entry into *K. pneumoniae* (33) and, thus circumvent the prospect of phage-resistance acquired by decreased expression of porins (43) and collateral increases in drug-resistant phenotype in the infection (44), we constructed as bait a strain that has no OmpK36. This Δ*ompK36* mutant strain of *K. pneumoniae* B5055 was constructed by “gene gorging” as previously described (45, 46) utilizing the donor and helper plasmids described in Supplementary Table 10.

### Phage isolation and infection of Klebsiella

Water samples were collected from catchment locations along the Merri Creek in Melbourne, Australia (Reservoir, postcode 3073, yielded MMNM, and Pascoe Vale, postcode 3044, yielded MMBB). Samples were centrifuged at 10,000× *g* for 10 minutes and filtered through a 0.45 μm cut-off filter. Water samples (45 mL) were subsequently mixed with 5 mL of 10× concentrated Luria-Bertani (LB) media and 1 mL of a *K. pneumoniae* B5055 Δ*ompK36* overnight culture and grown for a further 16 hours at 37°C. Cellular debris were pelleted by centrifugation at 10,000 × *g* for 10 minutes and the resulting supernatant was passed through a 0.45 μm filter. To monitor phage activity, 20 μL of the supernatant was then spotted onto LB agar plates containing a top layer of soft agar (4 mL LB and 0.35% (w/v) agar) and 200 μL of bacterial culture and incubated overnight at 37°C.

For liquid infections, the filtered supernatant was serially diluted with SM buffer (100 mM NaCl, 8 mM MgSO_4_, 10 mM Tris pH 7.5) and added to 200 μL of *K. pneumoniae* B5055 Δ*ompK36.* Cultures were incubated for 20 minutes at 37°C to allow phage adsorption and were then added to soft agar and poured using the double overlay method. Plaques with distinct morphologies were isolated from the top agar, serially diluted in SM buffer and incubated with the bacterial host as described above. This was repeated 5 times to obtain pure phage stocks.

### Phage amplification and purification

For large -amplification of the phages MMNM and MMBB, infections were performed using 14 cm petri dishes with 60 μL of phage preparation (10^−4^ dilution) added to 500 μL of an overnight culture and incubated for 20 minutes at 37°C. Ten millilitres of soft agar was then added to the culture and poured using the double agar layer method and incubated overnight at 37°C. Ten millilitres of SM buffer were added to each plate and incubated at room temperature for 10 minutes. The soft agar layer was scraped off using a disposable spreader and chloroform was subsequently added (1 mL/100 mL) to lyse bacterial cells to release the phages. The sample was then subject to vigorous shaking, before the agar and bacterial cell debris were removed by centrifugation at 11,000 × *g* for 40 minutes (4°C). The supernatant containing the phages was collected and DNase (1 μg/mL) and RNase (1 μg/mL) were subsequently added to the supernatant and incubated for 30 minutes at 4°C. NaCl (1 M final concentration) was added and incubated at 4°C for 1 hour with gentle mixing. Phages were precipitated from the media by adding PEG 8000 (10% final concentration) and incubated at 4°C overnight. Precipitated phage particles were collected by centrifugation at 11,000 × *g* for 20 minutes at 4°C and resuspended in SM buffer (1.6 mL/100 mL of precipitated supernatant). An equal volume of chloroform was added to the resuspended phage suspension to remove residual PEG and cell debris and vortexed for 30 seconds. The organic and aqueous phases were separated by centrifugation at 3,000 × *g* for 15 minutes at 4°C.

For purification on caesium chloride (CsCl) gradients, the aqueous phase containing the phages was removed and added to CsCl (0.5 g/mL of bacteriophage suspension) and mixed gently to dissolve the CsCl. The suspension was layered onto a discontinuous CsCl gradient (2 mL of 1.70 g/mL, 1.5 mL of 1.50 g/mL and 1.5 mL of 1.45 g/mL in SM buffer) in a Beckman SW41 centrifuge tube. Gradients were centrifuged at 22,000 rpm for 2 hours (4°C). Phage particles were collected from the gradient by piercing the side of the centrifuge tube with a syringe and removing the visible band in the gradient. Residual nucleic acid was removed from the phage preparation using floatation gradient centrifugation. Equal volumes of phage suspension (500 μL) and 7.2 M CsCl SM buffer were mixed and added to the bottom of a Beckman SW41 centrifuge tube. CsCl solutions (3 mL of 5 M and 7.5 mL of 3 M) were overlaid on top of the phage sample and centrifuged at 22,000 rpm for 2 hours (4°C). Phage particles were collected (~500 μL) using a syringe as described above. CsCl was dialysed out of the phage stock twice with 2 L of SM buffer overnight at 4 °C.

### Phage growth

One-step growth curve experiments were performed on *K. pneumoniae* as previously described (29). Mid-log-phase cultures were adjusted to an optical density at 600 nm (OD_600_) of 0.5, pelleted, and suspended in 0.1 volume of SM buffer. Phage lysate was subsequently added at a multiplicity of infection (MOI) of 0.01 and was allowed to adsorb for 10 minutes at 37°C. Following centrifugation at 12,000 × *g* for 4 minutes, the pellet was washed twice with SM buffer, resuspended with 30 mL of fresh LB broth, and incubated at 37°C. Samples were collected at 10-minute intervals for 120 minutes and titrated to determine PFU/mL. Growth experiments were performed in biological triplicates.

### Electron microscopy

From the CsCl-purifications, phage preparations (4 μL) were added to freshly glow-discharged CF200-Cu Carbon Support Film 200 Mesh Copper grids (ProScieTech) for 30 seconds. The sample was blotted from the grid using Whatman filter paper and samples were subsequently stained with 4 μL of Nano W Methylamine Tungstate (Nanoprobes) for 30 seconds and blotted again. Grids were imaged using a 120keV Tecnai Spirit G2 transmission electron microscope (Tecnai).

### Genomic DNA extraction, sequencing and annotation

Phage genomic DNA was isolated and samples were sequenced as 2× 250bp paired-end reads using Illumina MiSeq (36). The obtained reads were trimmed using Trimmomatic (47) and *de novo* assemblies of each genome were made using Burrows-Wheeler aligner (48) and Spades (49). The genomes were annotated using Prokka (50). The consensus sequences were then screened against the GenBank database using BLAST (https://blast.ncbi.nlm.nih.gov/Blast.cgi), date 29 April 2020. The genome data is available at Genbank with Accession ID: Klebsiella_phage_MMNM (MT894004) and Klebsiella_phage_MMBB (MT894005).

### Comparative genome analyses and BLAST

Proteomic trees were constructed using nucleotide genome sequences using the double stranded (ds) DNA nucleic acid type and Prokaryote host category database from ViPTree v1.9 (51) which also included a list of curated phage genomes (Supplementary Table 11). Refined trees were regenerated to analyse the phylogeny of either *Myoviridae* or *Siphoviridae* that infect *Gammaproteobacteria*. Each predicted open reading frame was analysed using BLASTP (https://blast.ncbi.nlm.nih.gov/Blast.cgi), Pfam HMMER (https://www.ebi.ac.uk/Tools/hmmer/) and HHpred (https://toolkit.tuebingen.mpg.de/) using the default settings.

A BLAST-based predictor was implemented during the evaluation of STEP^3^. It ran using blast-2.2.26+ For a query protein, the BLAST-based predictor will predict it to be positive if there is a BLAST hit against the training positive samples with a specified E-value. The E-value was set to 0.01 in this study, optimized on the independent dataset with a range of: 0.001, 0.01, 0.1, 1, and 10.

### Mass spectrometry

Each CsCl purified phage sample was solubilized in sodium dodecyl sulfate (SDS) lysis buffer (4% SDS, 100 mM HEPES pH8.5) and sonicated to assist protein extraction. The protein concentration was determined using a BCA kit (Thermo Scientific). SDS was removed according to previous work (52) and the proteins were proteolytically digested with trypsin (Promega) and purified using OMIX C18 Mini-Bed tips (Agilent Technologies) prior to LC-MS/MS analysis. Using a Dionex UltiMate 3000 RSLCnano system equipped with a Dionex UltiMate 3000 RS autosampler, an Acclaim PepMap RSLC analytical column (75 μm × 50 cm, nanoViper, C18, 2 μm, 100Å; Thermo Scientific) and an Acclaim PepMap 100 trap column (100 μm x 2 cm, nanoViper, C18, 5 μm, 100Å; Thermo Scientific), the tryptic peptides were separated by increasing concentrations of 80% acetonitrile / 0.1% formic acid at a flow of 250 nL/min for 120 minutes and analyzed with a QExactive Plus mass spectrometer (Thermo Scientific) using in-house optimized parameters to maximize the number of peptide identifications. To obtain peptide sequence information, the raw files were searched with Byonic v3.0.0 (ProteinMetrics) against the *K. pneumoniae* B5055 GenBank file FO834906 that was appended with the phage protein sequences. Only proteins falling within a false discovery rate (FDR) of 1% based on a decoy database were considered for further analysis.

### Raw data availability

The mass spectrometry proteomics data have been deposited to the ProteomeXchange Consortium via the PRIDE (53) partner repository with the dataset identifier PXD020607. **Username**, **Password**: ggYKM6wi

### Homology Modelling

Structural homologues were selected by querying the MMBB_78 sequence via the BLASTp webserver against the Protein Databank (PDB). In addition, this same sequence was probed using the Phyre2 software suite to identify local homology (54). Residues 186-872 of MMBB_78 were modelled against the enzymatic domain of the bacteriophage CBA120 tail-spike protein (PDB ID: 5W6P (55)). MODELLER v9.19 (56) was used with custom in-house scripts to generate 1000 potential models. These models were validated and sorted by their Discrete Optimised Protein Energy (DOPE) score, followed by visual inspection. An additional atomic model was calculated by the predictive software GalaxyTBM using the full length MMBB_78 sequence, as part of the GalaxyWEB (57) software suite.

### Construction of STEP^3^

#### Dataset construction

481 phage virion proteins were collected from the UniProt database with the “reviewed” tag and from the NCBI database following extensive literature searches. Redundant sequences were removed using the CD-HIT program (58) at a cut-off threshold of 0.4. As a result, 339 virion proteins with less than 40% sequence similarity were obtained. These proteins were further divided into two parts as positive samples: 243 in the training dataset and 96 in the independent dataset. For negative samples, 694 and 96 phage non-virion proteins were collected from UniProt to make up the training and independent datasets, respectively. Finally, a training dataset (243 positive samples and 694 negative samples) and an independent dataset (96 positive samples and 96 negative samples) were obtained, where each had less than 40% sequence similarity against each other. The two newly sequenced phage genomes MMNM and MMBB in this study were used to validate the prediction capability of STEP^3^ in practical scenarios.

#### PSSM generation

PSSM is a L*20 matrix, where L is the length of its original protein sequence and 20 is the number of amino acids. The (i, j)-th element (1<=i<=L, 1<=j<=20) in a PSSM corresponds to the probability of j-th amino acid to appear in the i-th position of its protein sequence. To generate a PSSM, blast-2.2.26 resource (ftp://ftp.ncbi.nlm.nih.gov/blast/executables) was used to search the protein sequence against the UniRef50 dataset (https://www.uniprot.org/help/uniref) with an E-value of 0.001 and the iteration of 3.

#### Feature encoding

Instead of extracting features directly from the protein sequences, evolutionary features mine patterns from a more informative profile in the format of PSSM. Five types of evolutionary features were generated using the POSSUM toolkit (59), including AAC-PSSM (60), PSSM-composition (61), DPC-PSSM (60), AADP-PSSM (60), and MEDP (62). For a given PSSM, their calculations are briefly described as follows: 1) AAC-PSSM generates a 20-dimensional vector through summing up and averaging all rows of the PSSM (60). 2) PSSM-composition further divides PSSM rows into 20 groups according to their corresponding amino acids in the original protein sequence (61). The rows in each group are summed up and normalized, and as a result the PSSM are transformed into a 20*20 matrix. Converting this matrix into a vector by row, PSSM-composition finally generates a 400-dimensional vector. 3) DPC-PSSM generates a 400-dimensional vector (*y*_l,l_,…, *y*_l,20_, *y*_2,l_,…, *y*_2,20_,…, *y*_20,l_,…, *y*_20,20_)^*T*^ through taking into account the local sequence-order effect (60). Among the vector, *y_i,j_* can be calculated by 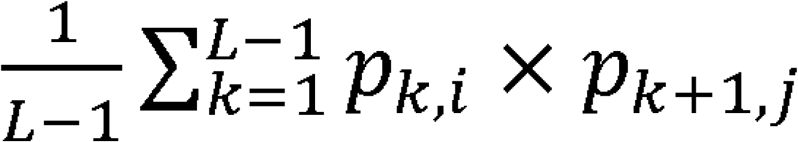 where i and j are between 1 and 20, and *p_k,i_* denotes the (k,i)-th element in PSSM. 4) AADP-PSSM combines AAC-PSSM and DPC-PSSM (60) as a 420-dimensional vector. 5) Likewise, MEDP generates a 420-dimensional vector through combining another two features EEDP and EDP (62). Among them, EEDP generates a 400-dimensional vector similarly to DPC-PSSM but using different transformation methodologies. EDP further sums up and averages all rows of the EEDP matrix to generate a 20-dimensional vector.

Additionally, four commonly used features were additionally implemented for comparison purpose, including the amino acid composition (AAC), dipeptide composition (DPC), QSOrder (63) and PAAC (64). AAC and DPC count the frequencies of residues and dipeptides in a protein sequence, respectively. QSOrder and PAAC extract features from a protein sequence as well, incorporating the physicochemical properties of its individual amino acids. Among them, QSOrder adopts Schneider-Wrede physicochemical distance matrix (65) and Grantham’s distance matrix (66), while PAAC takes hydrophobicity value from Tanford (67) and from Hopp and Woods (68), as well as amino acid side chain.

#### Model training on imbalanced data

Our imbalanced training dataset is to reflect the fact that the number of virion proteins is usually smaller than that of the non-virion proteins in a phage isolate. We combined all of the virion proteins with the same number of randomly selected non-virion proteins to generate a new balanced subset. This procedure was repeated five times, to generate five balanced subsets. For each feature, five individual models were trained based on five balanced subsets, and their prediction scores were averaged to obtain an ensemble model as the baseline model. Support vector machine (SVM) with a radial basis function kernel was used to train each model, implemented by the e1071 package (https://CRAN.R-project.org/package=e1071) in the R language (https://www.r-project.org/). The two parameters of SVM, including the Cost and Gamma, were optimized by a grid search between 2^−10^ and 2^10^ with a step of 2^1^ using the same R package.

#### Model integration

Training a model with each of features and then integrating them as an ensemble model usually have better and more robust performance, when compared with simply training a model with all features (69). Accordingly, the five baseline models (corresponding to five evolutionary features) were further integrated as the final ensemble model of STEP^3^ through averaging their prediction scores (Fig 1a).

#### Performance evaluation

The STEP^3^ predictor was extensively validated, with the baseline models and existing state-of-the-art tools on the 5-fold cross-validation and independent tests. Five performance metrics were used, including sensitivity (SN), specificity (SP), accuracy (ACC), F-value and Matthews correlation coefficient (MCC) (70). For each model, 5-fold cross-validation tests were conducted 5 times based on the 5 balanced training datasets, and then the performance metrics were averaged as the final performance result. The other tools compared to STEP^3^ were iVIREONS (https://vdm.sdsu.edu/ivireons), PVPred (http://lin-group.cn/server/PVPred), PVP-SVM (http://www.thegleelab.org/PVP-SVM/PVP-SVM.html), PVPred-SCM (http://camt.pythonanywhere.com/PVPred-SCM) and Pred-BVP-Unb (21). With no available tool for Pred-BVP-Unb, we developed one based on our training dataset by strictly following its methods, including its synthetic minority oversampling technique (SMOTE) to cope with the imbalance dataset, feature encodings, feature selection (a more geneneralized method GainRatio used) and the same grid search for parameter optimization. The prediction threshold for Pred-BVP-Unb is a standard cut-off of 0.5, which is the same as STEP^3^.

#### Sever construction and usage

The STEP^3^ server contains a client web interface and a server backend. The client web interface was implemented by the JAVA (https://www.java.com/) server development suite, JSP, CSS, jQuery (https://jquery.com/), Bootstrap (https://bootstrapdocs.com/) and their extension packages. The server backend was used by the Perl CGI (https://metacpan.org/pod/CGI). For visualization purposes, the blast 2.8.1+ (ftp://ftp.ncbi.nlm.nih.gov/blast/executables/blast+/2.8.1/) was used to search each predicted virion protein against known virion proteins to generate sequence similarities, which was visualized by BlasterJS (71). The MAFFT v7.271 (https://mafft.cbrc.jp/alignment/software/) was used to generate multiple alignment results between each predicted virion protein and known virion proteins, which was visualized by jsPhyloSVG (72). The all-against-all BLAST (version blast-2.2.26) was used to generate the sequence similarity network, visualized by ECharts (https://echarts.apache.org/). A queuing system was implemented using the Gearman framework (http://gearman.org/) to store the jobs the client deposits and dispatch them to idle threads maintained in the server backend. In this way, it links the two parts of STEP^3^, but decouples the prompt-response required in a client web interface and the time-consuming server backend for better user experience. To use the STEP^3^ server, users submit their protein sequences in FASTA format, and obtain a unique link to track the prediction progress or obtain the results once finished. In default mode, i.e. ‘For normal use’, the known virion proteins were marked with ‘exp.’ with an external link to the UniProt or NCBI database, while the predicted virion proteins were marked with ‘pred.’ with detailed annotations and options for visualization. Through interactive visualization, users could tentatively annotate the putative virion proteins with their potential subtype or functions, based on the sequence similarity or phylogenetic analysis considerations. For users who want to benchmark the STEP^3^ server, a ‘For benchmarking test’ option is available to obtain prediction scores for all their sequences.

## ACKNOWLEDGEMENTS

We acknowledge that this project was conducted on the traditional homelands of the Wurundjeri Woi wurrung people, with the phages isolated from waters of the Merri Creek, Melbourne, Australia. The Centre to Impact AMR would like to acknowledge and thank Wurundjeri Woi wurrung Elder, Aunty Gail Smith, who named the phages in this study in Woi wurrung language. Merri-merri-uth nyilam marra-natj (MMNM) and Merri-merri baany-a bundha-natj (MMBB) translate as “Dangerous Merri lurker” and “Merri water biter”, respectively, in English. Our future work in this field will be pursued according to a Memorandum of Understanding (MoU) between the Monash Centre to Impact AMR and the Wurundjeri Woi wurrung Cultural Heritage Aboriginal Corporation (https://www.wurundjeri.com.au/) the peak body representing the Wurundjeri Woi wurung people. The MoU recognizes the Wurundjeri Woi wurrung as the sovereign First People of their Country with distinct rights, and will ensure the equitable sharing of resources including any commercial benefits realized from the development of Wurundjeri Woi wurrung resources. We acknowledge Jordan Smith and Karmen Jobling of the Wurundjeri Woi wurrung Cultural Heritage Aboriginal Corporation’s Water Unit for their stewardship in shaping the MoU between Wurundjeri Woi wurrung Cultural Heritage Aboriginal Corporation and Monash Centre to Impact AMR. We are grateful to Professor Richard Strugnell, Department of Microbiology and Immunology, University of Melbourne for access to his collection of *Klebsiella* isolates. W.D. was a visiting MSc student at Monash University, supported by the study abroad program for graduate student of Guilin University of Electronic Technology (GDYX2019010). Research was supported by a seed grant from the Monash-Warwick Alliance (to T.L., E.J. and S.McG) and the initial phase of the project was supported by the Australian Research Council (FL130100038).

## AUTHOR CONTRIBUTIONS

TTY, MEW and JJW performed biological experiments. WD and YZ performed computational experiments. DW, RSB and SMcG performed structural calculations and modelling, and RSB performed electron microscopy. EJ, JJB, AR, C.J.S., CH and RS analyzed data. JW, RAD and TL supervised project, analyzed data and wrote the paper. All authors contributed critical evaluation to the final version of the manuscript.

## COMPETING INTERESTS

The authors declare no competing interests.

## SUPPLEMENT

**Supplementary Figure 1. Sequence analysis and data visualization of the major capsid protein from λ phage**.

**Supplementary Figure 2. Comparative genome analysis of *Klebsiella* phage MMBB**.

**Supplementary Figure 3. Structure informed analysis of *Klebsiella* phage MMBB virion components**.

**Supplementary Table 1. Prediction performance of STEP^3^ and baseline models on the 5-fold cross-validation test**.

**Supplementary Table 2. Prediction performance of STEP^3^ and baseline models on the on the independent test**.

**Supplementary Table 3. Prediction performance of STEP^3^, other available predictors and the BLAST-based baseline predictor on the independent dataset**.

**Supplementary Table 4. Annotation of *Klebsiella* phage MMNM genome**.

**Supplementary Table 5. Annotation of *Klebsiella* phage MMBB genome**.

**Supplementary Table 6. Mass spectrometry and STEP^3^ analysis of *Klebsiella* phage MMNM virions**.

**Supplementary Table 7. Detailed prediction of STEP^3^, other available predictors and the BLAST-based baseline predictor on the phage *Klebsiella* phage MMNM**.

**Supplementary Table 8. Detailed prediction of STEP^3^, other available predictors and the BLAST-based baseline predictor on the phage *Klebsiella* phage MMBB**.

**Supplementary Table 9. Mass spectrometry and STEP^3^ analysis of *Klebsiella* phage MMBB virions**.

**Supplementary Table 10. Strains, plasmids and primers used in this study**.

**Supplementary Table 11. Genomes of phages used for tree analysis.**

## Notes

http://step3.erc.monash.edu/

### Competing Interest Statement

The authors have declared no competing interest.

