## Supplemental data for "The component parts of bacteriophage virions accurately defined by a machine-learning approach built on evolutionary features"

**Supplementary Figure 1. Sequence analysis and data visualization of the major capsid protein from λ phage.**

**Supplementary Figure 2. Comparative genome analysis of *Klebsiella* phage MMBB.**

**Supplementary Figure 3. Structure informed analysis of *Klebsiella* phage MMBB virion components.**

**Supplementary Table 1. Prediction performance of $STEP^3$ and baseline models on the 5-fold cross-validation test.**

**Supplementary Table 2. Prediction performance of $STEP^3$ and baseline models on the on the independent test.**

**Supplementary Table 3. Prediction performance of $STEP^3$, other available predictors and the BLAST-based baseline predictor on the independent dataset.**

**Supplementary Table 4. Annotation of *Klebsiella* phage MMNM genome.**

**Supplementary Table 5. Annotation of *Klebsiella* phage MMBB genome.**

**Supplementary Table 6. Mass spectrometry and $STEP^3$ analysis of *Klebsiella* phage MMNM virions.**

**Supplementary Table 7. Detailed prediction of $STEP^3$, other available predictors and the BLAST-based baseline predictor on the phage *Klebsiella* phage MMNM.**

**Supplementary Table 8. Detailed prediction of $STEP^3$, other available predictors and the BLAST-based baseline predictor on the phage *Klebsiella* phage MMBB.**

**Supplementary Table 9. Mass spectrometry and $STEP^3$ analysis of *Klebsiella* phage MMBB virions.**

**Supplementary Table 10. Strains, plasmids and primers used in this study.**

**Supplementary Table 11. Genomes of phages used for tree analysis.**

**a**

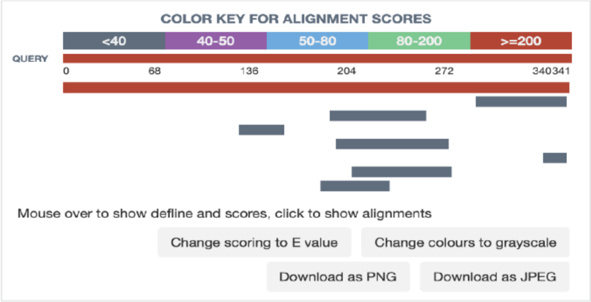

| Description | Max score | Total score | Query cover | E value | Identities |
|---|---|---|---|---|---|
| P68650 Major capsid protein OS=Enterobacteria phage P21 OX=10711 GN=E PE=3 SV=1 | 431 | 431.0 | 99% | 3e-154 | 62% |
| Q01076 Internal virion protein gp20 OS=Salmonella phage P22 OX=10754 GN=20 PE=1 SV=2 | 26.2 | 26.2 | 18% | 0.39 | 27% |
| P10928 Baseplate wedge protein gp10 OS=Enterobacteria phage T4 OX=10665 GN=10 PE=1 SV=1 | 25.4 | 25.4 | 19% | 0.77 | 30% |
| P16009 Baseplate central spike complex protein gp5 OS=Enterobacteria phage T4 OX=10665 GN=5 PE=1 SV=2 | 22.7 | 22.7 | 9% | 4.8 | 29% |
| Q4Z9F2 Tail sheath protein OS=Staphylococcus phage Twort (strain DSM 17442) OX=1283338 PE=3 SV=1 | 22.7 | 22.7 | 22% | 5.3 | 30% |
| P13390 L-shaped tail fiber protein pb1 OS=Escherichia phage T5 OX=10726 GN=ltf PE=1 SV=3 | 22.3 | 22.3 | 4% | 7.0 | 53% |
| P25129 Attachment protein G3P OS=Pseudomonas phage Pf1 OX=2011081 GN=III PE=3 SV=1 | 21.9 | 21.9 | 19% | 7.8 | 28% |
| Q6QGF0 Probable central straight fiber OS=Escherichia phage T5 OX=10726 GN=D17 PE=2 SV=1 | 21.9 | 21.9 | 13% | 10.0 | 36% |

Download as CSV   Download as PNG   Download as JPEG

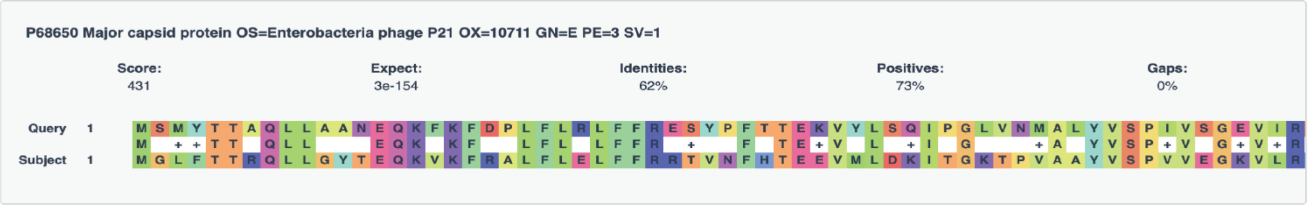

**Supplementary Figure 1. Sequence analysis and data visualization of the major capsid protein from λ phage.** (a) Conserved aspects of domain architecture based on gpE, the major capsid protein of λ phage from BLASTp analysis, as documented under the “Sequence similarity” section of the data visualization section in STEP[3] [continued over-page].

**b**

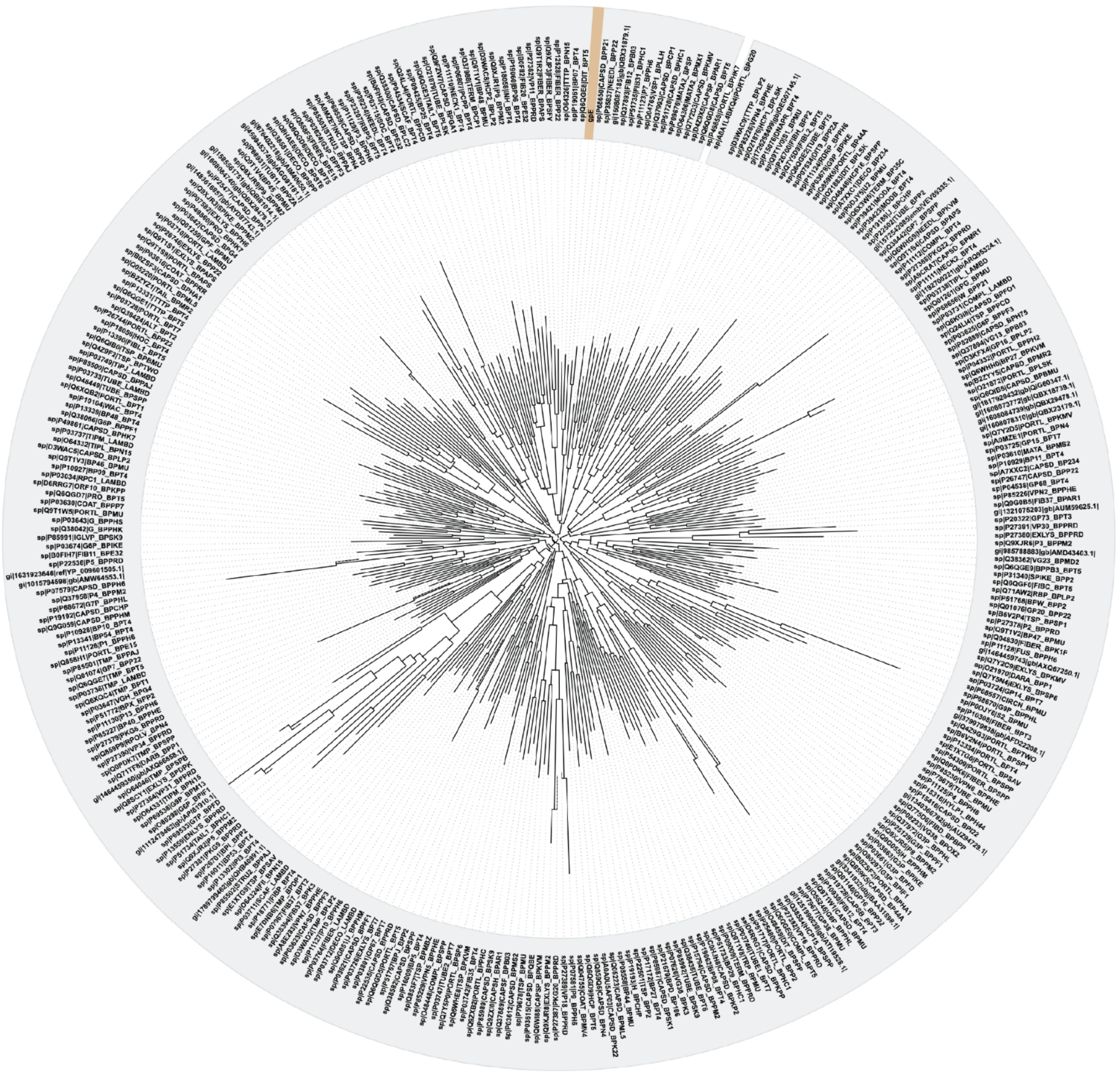

**Supplementary Figure 1. Sequence analysis and data visualization of the major capsid protein from λ phage.** (b) Protein similarity relationships for λ phage gpE generated and visualized in the "Phylogenetic tree" of STEP[3] [continued over-page].

c

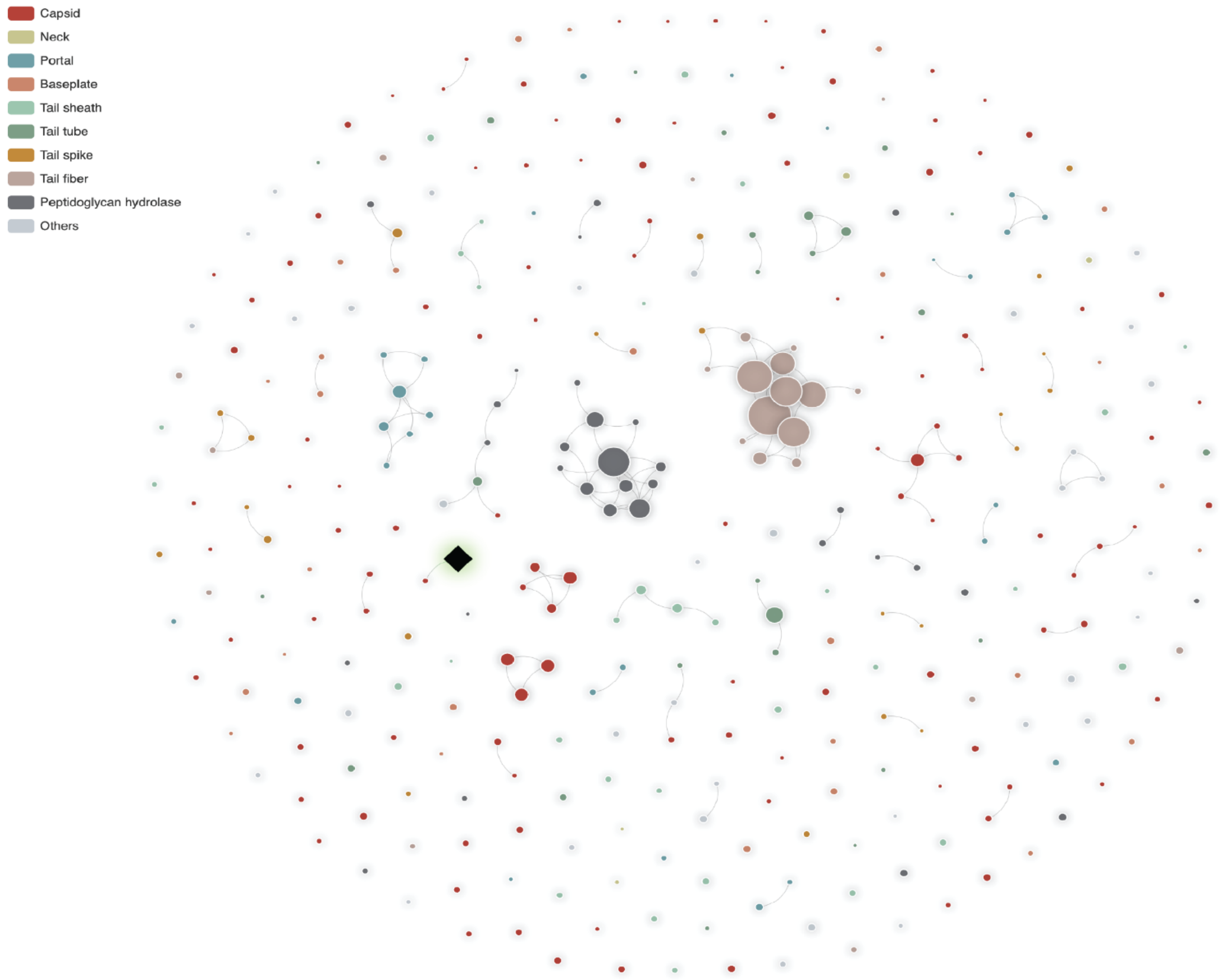

**Supplementary Figure 1. Sequence analysis and data visualization of the major capsid protein from λ phage.** (c) Relationships for λ phage gpE (♦) as documented under the "Homology network visualization 1" section of the data visualization section in STEP[3] [continued over-page].

d

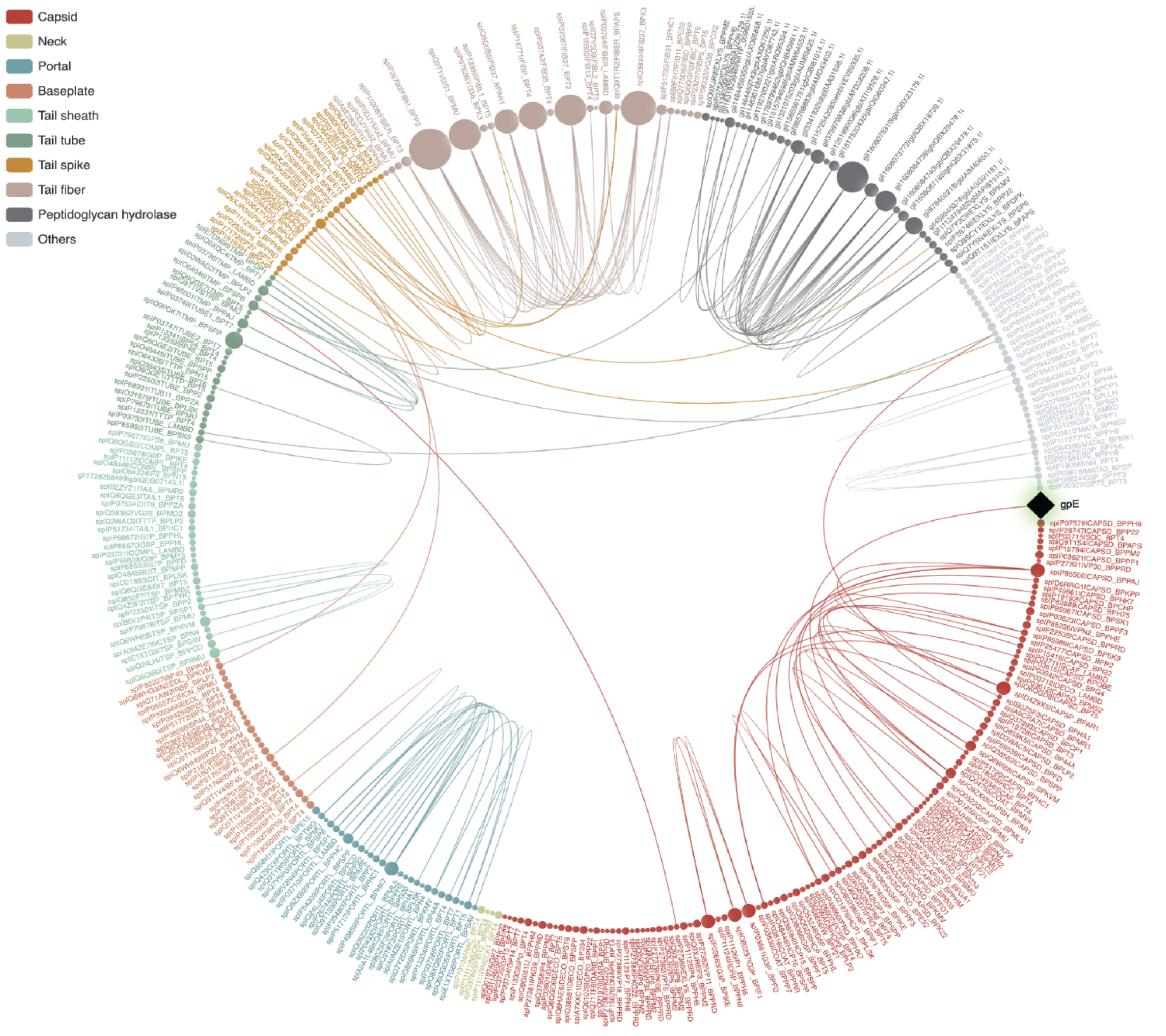

**Supplementary Figure 1. Sequence analysis and data visualization of the major capsid protein from λ phage.** (d) Relationships for λ phage gpE (♦) visualized with the "Homology network visualization 2" section of STEP[3].

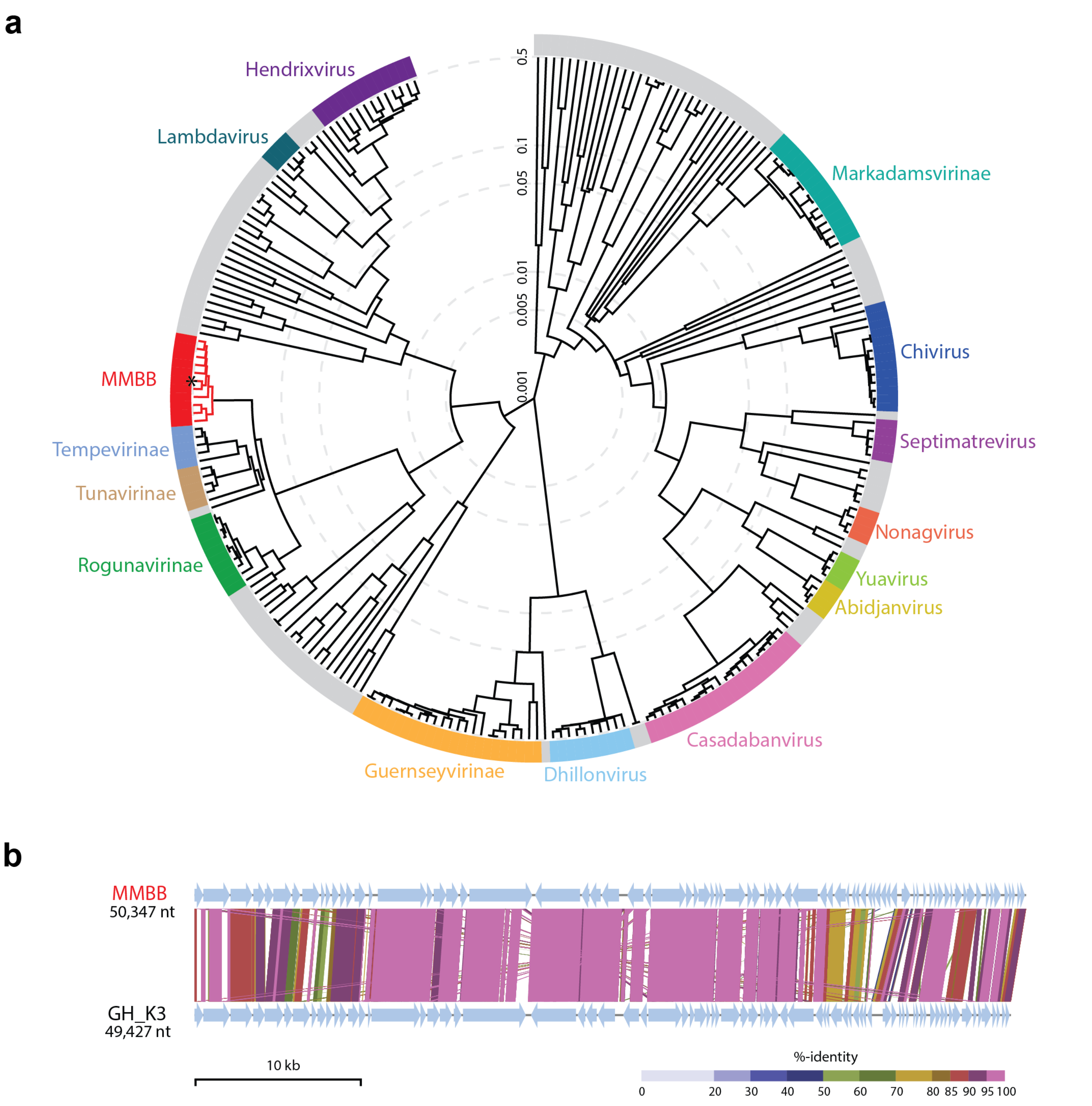

**Supplementary Figure 2. Comparative genome analysis of *Klebsiella* phage MMBB.** (a) Proteomic tree analysis of *Siphoviridae*-like phages that infect *Gammaproteobacteria* generated using ViPTree[1]. The branch lengths represent genomic similarity based on normalised tBLASTx scores plotted on a logarithmic scale. The tree was constructed using sequences from the default ViPTree dataset and other selected phage genomes listed in Table S11. Viral subfamilies or genera are highlighted in the coloured bars. Gray bars represent phages that belong to groups that are currently unclassified. Members of the *Webervirus* group, including *Klebsiella* phage MMBB (*), are highlighted in red. (b) Whole genome alignment of phages *Klebsiella* phage MMBB and GH_K3 created using ViPTree. Each genome has been orientated to start with the putative small terminase subunit gene. For panel b. the sequences are linked by colored bars highlighting sequence identity values as shown in the key at the foot of the Figure.

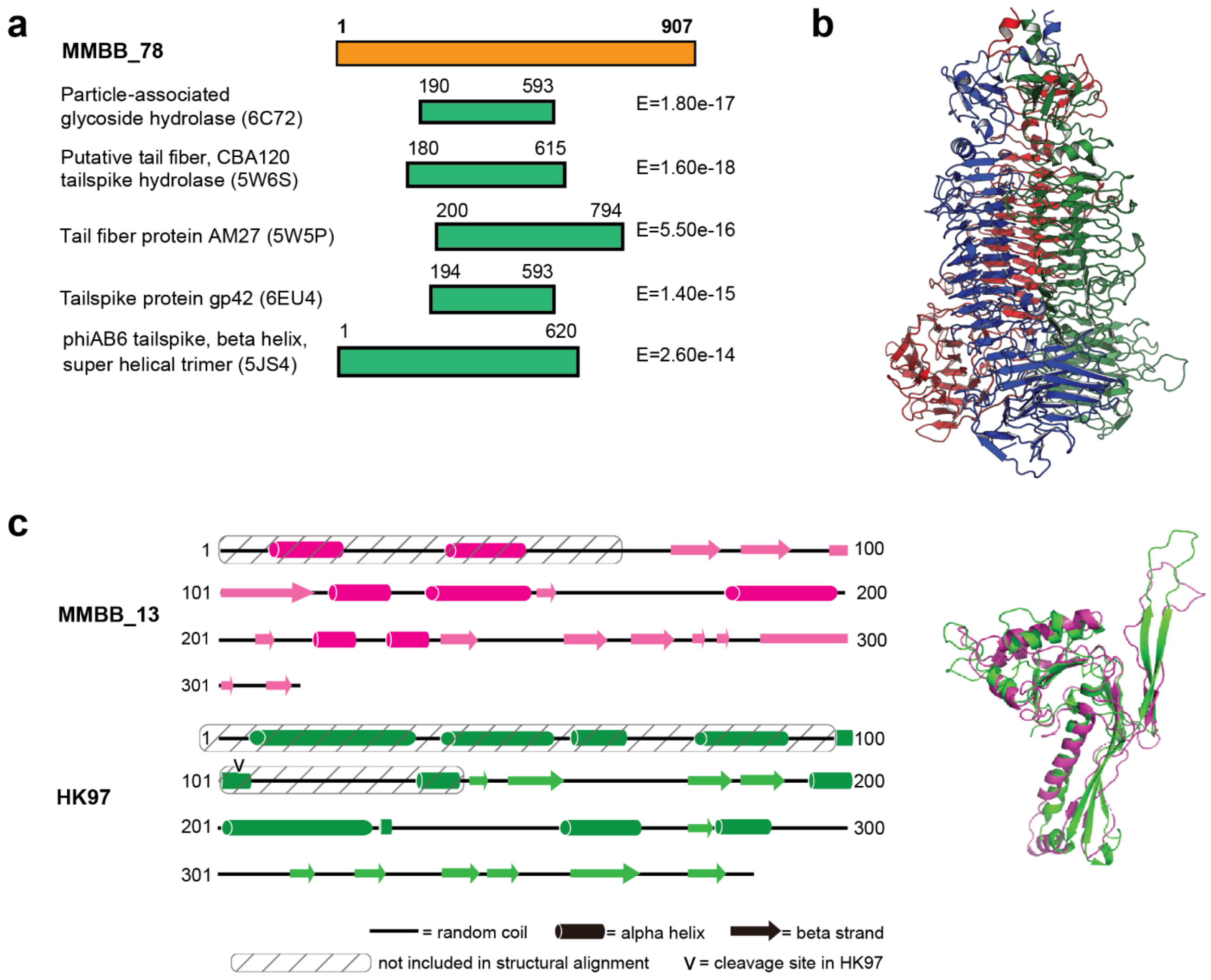

**Supplementary Figure 3. Structure informed analysis of *Klebsiella* phage MMBB virion components.** (a) Representation of the 907 residues in MMBB_78, and the and the segments that show structural similarity to proteins of known structure. The PDB coordinates for each of these proteins are shown. (b) Structural model of the putative depolymerase MMBB_78. Analysis with HHpred for remote protein homology detection and structure prediction is shown, consistent with a pectate lyase-like right-handed β-helix repeat structure as depicted, structurally homologous to the CBA120 tail-spike protein (PDB ID: 5W6P) (c) While both MMBB_13 and MMBB_15 were annotated as "major capsid protein", Psipred[74] secondary structure predictions showed only MMBB_13 (pink) as conforming to the HK97 fold observed in all major capsid proteins (green). 3D-structure predictions demonstrate this HK97 structural similarity (pink, Phyre2[61] model of MMBB_13) when aligned with the structure of the phage HK97 head II protein (PDB ID 2FT1).

**Supplementary Table 1. Prediction performance of STEP$^3$ and baseline models using the 5-fold cross-validation test**

| Encoding | Sensitivity (SN) | Specificity (SP) | Accuracy (ACC) | F-value | Matthews correlation coefficient (MCC) |
|---|---|---|---|---|---|
| AAC | 0.753±0.036 | 0.782±0.040 | 0.767±0.019 | 0.762±0.020 | 0.537±0.037 |
| DPC | 0.754±0.017 | 0.793±0.023 | 0.774±0.013 | 0.768±0.013 | 0.548±0.028 |
| QSOrder | 0.770±0.018 | 0.793±0.022 | 0.780±0.020 | 0.776±0.020 | 0.565±0.038 |
| PAAC | 0.760±0.019 | 0.793±0.048 | 0.778±0.023 | 0.772±0.019 | 0.558±0.043 |
| AAC-PSSM | **0.817±0.017** | 0.858±0.022 | **0.837±0.017** | **0.832±0.016** | **0.675±0.035** |
| PSSM-composition | 0.797±0.019 | 0.828±0.023 | 0.812±0.019 | 0.809±0.019 | 0.625±0.037 |
| DPC-PSSM | 0.778±0.010 | 0.844±0.016 | 0.811±0.009 | 0.803±0.009 | 0.622±0.019 |
| AADP-PSSM | 0.788±0.008 | 0.838±0.014 | 0.812±0.009 | 0.806±0.009 | 0.625±0.020 |
| MEDP | 0.807±0.025 | 0.825±0.024 | 0.816±0.020 | 0.813±0.021 | 0.631±0.041 |
| Step$^3$ | 0.803±0.025 | **0.864±0.015** | 0.833±0.017 | 0.826±0.019 | 0.667±0.034 |

*Note*: Values are in the form of mean ± standard deviation. The best performance value for each metric across different models is highlighted in bold. Five performance metrics were used as indicated in the column headers. STEP$^3$ refers to the final ensemble model that integrated five evolutionary feature-based baseline models by averaging their prediction scores. This applies to all other results if not explicitly specified.

**Supplementary Table 2. Prediction performance of STEP$^3$ and baseline models using the independent test**

| Encoding | Sensitivity (SN) | Specificity (SP) | Accuracy (ACC) | F-value | Matthews correlation coefficient (MCC) |
|---|---|---|---|---|---|
| AAC | 0.875 | 0.812 | 0.844 | 0.848 | 0.689 |
| DPC | 0.812 | 0.875 | 0.844 | 0.839 | 0.689 |
| QSOrder | 0.844 | 0.844 | 0.844 | 0.844 | 0.688 |
| PAAC | 0.854 | 0.833 | 0.844 | 0.845 | 0.688 |
| AAC-PSSM | 0.885 | 0.854 | 0.87 | 0.872 | 0.74 |
| PSSM-composition | **0.906** | 0.844 | 0.875 | 0.879 | 0.751 |
| DPC-PSSM | 0.875 | 0.844 | 0.859 | 0.862 | 0.719 |
| AADP-PSSM | 0.896 | 0.865 | 0.88 | 0.882 | 0.761 |
| MEDP | 0.885 | 0.854 | 0.87 | 0.872 | 0.74 |
| STEP$^3$ | 0.896 | **0.885** | **0.891** | **0.891** | **0.781** |

*Note*: The best performance value for each metric across different models is highlighted in bold.

**Supplementary Table 3. Prediction performance of STEP[3], other available predictors and the BLAST-based baseline predictor on the independent dataset**

| **Model** | **Sensitivity (SN)** | **Specificity (SP)** | **Accuracy (ACC)** | **F-value** | **Matthews correlation coefficient (MCC)** |
|---|---|---|---|---|---|
| BLAST | 0.25 | **0.99** | 0.63 | 0.423 | 0.375 |
| iVIREONS | 0.781 | 0.698 | 0.74 | 0.75 | 0.481 |
| PVPred | 0.385 | 0.875 | 0.63 | 0.51 | 0.299 |
| PVP-SVM | 0.427 | 0.865 | 0.646 | 0.547 | 0.324 |
| Pred-BVP-Unb* | 0.615 | 0.917 | 0.766 | 0.724 | 0.557 |
| PVPred-SCM | 0.51 | 0.76 | 0.635 | 0.583 | 0.28 |
| STEP[3] | **0.896** | 0.885 | **0.891** | **0.891** | **0.781** |

*Note*: The best performance value for each metric across different predictors is highlighted in bold.
(*) Denotes that Pred-BVP-Unb is not publicly available so was recreated according to published methodology from Arif *et al.*[2].

**Supplementary Table 4. Annotation of *Klebsiella* phage MMNM genome.**

| ORF | start | end | BLASTp, HMMer and HHpred Description | BLASTp analysis | Protein sequence identity[b] | Accession no. |
|---|---|---|---|---|---|---|
| 1 | 468 | 1463 | Endonuclease | Putative Hef-like homing endonuclease [*Klebsiella* phage 1611E-K2-1] | 272/328 (83) | ATS92554 |
| 2 | 1496 | 1813 | Hypothetical protein | Hypothetical protein kpv52_20 [*Klebsiella* phage vB_KpnM_KpV52] | 94/105 (90) | YP_009597548 |
| 3 | 1875 | 2294 | Putative structural protein[a], DUF4054 | Hypothetical protein [Myoviridae sp.] | 124/139 (89) | QHJ80433 |
| 4 | 2304 | 2933 | Putative structural protein[a] | Hypothetical protein kpv79_21 [*Klebsiella* phage vB_KpnM_KpV79] | 191/209 (91) | YP_009615292 |
| 5 | 2958 | 3089 | Hypothetical protein | Hypothetical protein kpv79_22 [*Klebsiella* phage vB_KpnM_KpV79] | 41/43 (95) | YP_009615293 |
| 6 | 3086 | 3385 | Hypothetical protein | Hypothetical protein [*Klebsiella* phage JD001] | 97/98 (99) | YP_007392835 |
| 7 | 3975 | 4478 | Neck protein | Neck protein [*Klebsiella* phage vB_KpnM_FZ14] | 154/166 (93) | QCG76511 |
| 8 | 4487 | 4828 | Head-closure protein | Head-closure protein [*Klebsiella* phage vB_KpnM_FZ14] | 111/113 (98) | QCG76512 |
| 9 | 4815 | 5396 | Tail-completion protein | Tail-completion protein [*Klebsiella* phage vB_KpnM_FZ14] | 187/193 (97) | QCG76513 |
| 10 | 5439 | 6572 | Putative structural protein[a] | Hypothetical protein FZ14_34 [*Klebsiella* phage vB_KpnM_FZ14] | 361/377 (96) | QCG76514 |
| 11 | 6586 | 7002 | Putative structural protein[a] | Hypothetical protein kpv79_28 [*Klebsiella* phage vB_KpnM_KpV79] | 135/138 (98) | YP_009615299 |
| 12 | 7041 | 7757 | DNA-binding Protein | Putative DNA-binding protein [*Klebsiella* phage vB_KpnM_KpV52] | 222/238 (93) | YP_009597558 |
| 13 | 7754 | 7942 | DNA-binding protein | DNA-binding protein [*Klebsiella* phage vB_KpnM_FZ14] | 54/59 (92) | QCG76483 |
| 14 | 8025 | 8207 | Hypothetical protein | Hypothetical protein [*Klebsiella* phage vB_KpnM_IME346] | 134/138 (97) | QBZ68924 |
| 15 | 8272 | 8985 | DNA-binding protein | DNA-binding protein [*Klebsiella* phage vB_KpnM_KpV79] | 121/131 (92) | YP_009615303 |
| 16 | 9111 | 9485 | Putative structural protein[a] | Hypothetical protein kpv79_33 [*Klebsiella* phage vB_KpnM_KpV79] | 123/124 (99) | YP_009615304 |
| 17 | 9672 | 9968 | Putative structural protein[a] | Hypothetical protein kpv52_35 [*Klebsiella* phage vB_KpnM_KpV52] | 95/98 (97) | YP_009597563 |

| | | | | | | |
|---|---|---|---|---|---|---|
| 18 | 9968 | 11368 | Tape measure protein | Tape measure protein [*Klebsiella* phage vB_KpnM_IME346] | 426/477 (89) | QBZ68918 |
| 19 | 11371 | 12795 | Lytic transglycosylase | Putative lytic transglycosylase [*Klebsiella* phage vB_KpnM_15-38_KLPPOU148] | 456/474 (96) | QGZ13372 |
| 20 | 12792 | 13442 | Putative structural protein[a] | Hypothetical protein KLPPOU148_072 [*Klebsiella* phage vB_KpnM_15-38_KLPPOU148] | 213/216 (99) | QGZ13389 |
| 21 | 13732 | 14406 | Baseplate protein | Putative baseplate protein [*Klebsiella* phage vB_KpnM_IME346] | 222/224 (99) | QBZ68915 |
| 22 | 14406 | 14753 | Phospholipase | Phospholipase [*Klebsiella* phage vB_KpnM_FZ14] | 113/115 (98) | QCG76487 |
| 23 | 14746 | 15954 | Baseplate protein | Baseplate protein J-like protein [*Klebsiella* phage vB_KpnM_FZ14] | 394/402 (98) | QCG76488 |
| 24 | 15951 | 17957 | Tail fibre protein, hydrolase, endo-glycosidase | Putative tail fibre family protein [*Klebsiella* phage vB_KpnM_FZ14] | 645/668 (97) | QCO71663 |
| 25 | 17972 | 18145 | Putative structural protein[a] | Hypothetical protein kpv52_43 [*Klebsiella* phage vB_KpnM_KpV52] | 56/57 (98) | YP_009597571 |
| 26 | 18246 | 18959 | Coil containing Protein | Coil containing protein [*Klebsiella* phage vB_KpnM_FZ14] | 225/237 (95) | QCG76489 |
| 27 | 18963 | 19856 | Tail fibre protein | Putative tail fibre protein [*Klebsiella* phage vB_KpnM_KpV52] | 284/297 (96) | YP_009597573 |
| 28 | 19851 | 20087 | Outer membrane spanin subunit | Putative outer membrane spanin subunit [*Klebsiella* phage vB_KpnM_IME346] | 73/78 (94) | QBZ68906 |
| 29 | 20084 | 20410 | Inner membrane spanin subunit | Putative inner membrane spanin subunit [*Klebsiella* phage vB_KpnM_IME346] | 95/108 (88) | QBZ68905 |
| 30 | 20410 | 20976 | Endolysin | Putative endolysin [*Klebsiella* phage vB_KpnM_15-38_KLPPOU148] | 180/184 (98) | QGZ13395 |
| 31 | 21022 | 21138 | Hypothetical protein | Hypothetical protein kpv52_49 [*Klebsiella* phage vB_KpnM_KpV52] | 37/38 (97) | YP_009597577 |
| 32 | 21142 | 21564 | Hypothetical protein | Hypothetical protein [*Klebsiella* phage JD001] | 139/140 (99) | YP_007392880 |
| 33 | 21579 | 21860 | Putative structural protein[a] | Hypothetical protein [*Klebsiella* phage JD001] | 93/93 (100) | YP_007392879 |
| 34 | 21877 | 22218 | Putative DNA binding protein | Hypothetical protein kpv52_52 [*Klebsiella* phage vB_KpnM_KpV52] | 113/113 (100) | YP_009597580 |

| | | | | | | |
|---|---|---|---|---|---|---|
| 35 | 22215 | 24389 | DNA polymerase | Putative DNA polymerase [*Klebsiella* phage vB_KpnM_KpV52] | 695/724 (96) | YP_009597581 |
| 36 | 24437 | 24553 | Hypothetical protein | Hypothetical protein kpv79_56 [*Klebsiella* phage vB_KpnM_KpV79] | 35/36 (97) | YP_009615327 |
| 37 | 24604 | 25416 | Putative structural protein[a] | Hypothetical protein kpv52_56 [*Klebsiella* phage vB_KpnM_KpV52] | 266/270 (99) | YP_009597584 |
| 38 | 25503 | 26627 | Putative ATP dependant DNA helicase | Hypothetical protein [*Klebsiella* phage JD001] | 347/374 (93) | YP_007392872 |
| 39 | 26633 | 27490 | Putative structural protein[a] | Hypothetical protein kpv79_59 [*Klebsiella* phage vB_KpnM_KpV79] | 260/292 (89) | YP_009615330 |
| 40 | 27585 | 27860 | Hypothetical protein | Hypothetical protein [*Klebsiella* phage vB_KpnM_IME346] | 72/75 (96) | QBZ68968 |
| 41 | 27857 | 28027 | Hypothetical protein | Hypothetical protein [*Klebsiella* phage vB_KpnM_IME346] | 50/53 (94) | QBZ68967 |
| 42 | 28086 | 30785 | Helicase | Putative helicase [*Klebsiella* phage JD001] | 846/898 (94) | YP_007392869 |
| 43 | 30820 | 31290 | Putative nuclease | Hypothetical protein kpv79_65 [*Klebsiella* phage vB_KpnM_KpV79] | 149/156 (96) | YP_009615336 |
| 44 | 31353 | 31631 | TetR transcriptional regulator | Putative TetR transcriptional regulator [*Klebsiella* phage vB_KpnM_KpV52] | 86/92 (93) | YP_009597595 |
| 45 | 31654 | 32547 | Putative DNA directed RNA polymerase subunit | Hypothetical protein kpv79_67 [*Klebsiella* phage vB_KpnM_KpV79] | 106/140 (76) | YP_009615338 |
| 46 | 32755 | 35025 | DNA primase | DNA primase [*Klebsiella* phage vB_KpnM_KpV79] | 722/756 (96) | YP_009615339 |
| 47 | 35117 | 35290 | HAD superfamily hydrolase | HAD superfamily hydrolase [*Klebsiella* phage JD001] | 54/56 (96) | YP_007392864 |
| 48 | 35278 | 35481 | Hypothetical protein | Hypothetical protein kpv79_70 [*Klebsiella* phage vB_KpnM_KpV79] | 64/67 (96) | YP_009615341 |
| 49 | 35604 | 36728 | Putative DNA polymerase II small subunit | Hypothetical protein KLPPOU148_031 [*Klebsiella* phage vB_KpnM_15-38_KLPPOU148] | 359/374 (96) | QGZ13378 |
| 50 | 36725 | 37219 | Polynucleotide kinase | Putative polynucleotide kinase/phosphatase [*Klebsiella* phage VLC6] | 125/163 (77) | QJ152607 |
| 51 | 37216 | 37377 | Hypothetical protein | Hypothetical protein [Bacteriophage sp.] | 48/52 (92) | QHJ82135 |
| 52 | 37411 | 37905 | Hypothetical protein | Hypothetical protein [*Klebsiella* phage vB_KpnM_IME346] | 158/164 (96) | QBZ68954 |

| | | | | | | |
|---|---|---|---|---|---|---|
| 53 | 38490 | 38948 | Small subunit terminase | Small subunit terminase [*Klebsiella* phage vB_KpnM_IME346] | 151/152 (99) | QBZ68953 |
| 54 | 38938 | 40386 | Large subunit terminase | Large subunit terminase [*Klebsiella* phage vB_KpnM_FZ14] | 476/482 (99) | QCG76505 |
| 55 | 40389 | 41783 | Portal protein | Portal protein [Pantoea phage vB_PagM_AAM22] | 345/472 (73) | QDH45578 |
| 56 | 41799 | 42008 | Putative DNA directed RNA polymerase subunit | Hypothetical protein [Serratia sp. Nf2] | 39/63 (62) | WP_107227763 |
| 57 | 42005 | 42319 | Putative structural protein[a] | Hypothetical protein [Cronobacter dublinensis] | 38/67 (57) | WP_105739368 |
| 58 | 42377 | 43141 | Head morphogenesis protein | Head morphogenesis protein [*Klebsiella* phage vB_KpnM_KpV79] | 243/254 (96) | YP_009615278 |
| 59 | 43200 | 43397 | Outer membrane adhesin like protein | Outer membrane adhesin like protein [*Klebsiella* phage vB_KpnM_IME346] | 61/65 (94) | QBZ68946 |
| 60 | 43394 | 43543 | Glycosyltransferase | Putative glycosyltransferase [*Klebsiella* phage vB_KpnM_KpV79] | 44/49 (90) | YP_009615280 |
| 61 | 43614 | 43808 | Hypothetical protein | Hypothetical protein P_46 [Escherichia phage P AB-2017] | 51/63 (81) | AQN31984 |
| 62 | 43792 | 43989 | Putative structural protein[a] | Hypothetical protein kpv79_10 [*Klebsiella* phage vB_KpnM_KpV79] | 52/58 (90) | YP_009615281 |
| 63 | 43989 | 44174 | Hypothetical protein | Hypothetical protein [*Klebsiella* phage JD001] | 39/39 (100) | YP_007392847 |
| 64 | 44174 | 44395 | Hypothetical protein | Hypothetical protein kpv52_15 [*Klebsiella* phage vB_KpnM_KpV52] | 66/73 (90) | YP_009597543 |
| 65 | 44472 | 45566 | Coil containing protein | Coil containing protein [*Klebsiella* phage vB_KpnM_FZ14] | 348/364 (96) | QCG76509 |
| 66 | 45578 | 46066 | Putative capsid fibre protein | Hypothetical protein [*Klebsiella* phage JD001] | 148/162 (91) | YP_007392843 |
| 67 | 46069 | 47103 | Major capsid protein | Major capsid protein [*Klebsiella* phage vB_KpnM_KpV79] | 315/343 (92) | YP_009615287 |

*Note*: [a], proteins that were identified by mass spectrometry analysis of MMNM virions and were predicted to be a virion-associated protein by STEP[3] but had been previously annotated only as "hypothetical proteins". [b], Protein sequence identity is expressed as a ratio of identical residues/total residues and, in parentheses, as a % identity.

**Supplementary Table 5. Annotation of *Klebsiella* phage MMBB genome.**

| ORF | start | end | BLASTp, HMMer and HHpred Description | BLASTp analysis | Protein sequence identity[b] | Accession no. |
|---|---|---|---|---|---|---|
| 1 | 2295 | 2897 | Tail assembly protein | Putative tail assembly protein [*Klebsiella* phage TSK1] | 200/200 (100) | AXN53798 |
| 2 | 2872 | 3609 | Minor tail protein | Minor tail protein [*Klebsiella* phage NJS3] | 244/245 (99) | AXQ68096 |
| 3 | 3611 | 4363 | Minor tail protein L | Minor tail protein [*Klebsiella* phage MezzoGao] | 113/114 (99) | ASV44957 |
| 4 | 4433 | 4777 | Minor tail protein | Minor tail protein [*Klebsiella* phage MezzoGao] | 113/114 (99) | YP_00979211 0 |
| 5 | 4780 | 7749 | Tail length tape-measure protein | Tail length tape-measure protein [*Klebsiella* phage NJR15] | 974/989 (98) | AXQ68170 |
| 6 | 8100 | 8426 | Tail assembly protein | Tail assembly chaperone [*Klebsiella* phage Shelby] | 108/108 (100) | QEG07301 |
| 7 | 8503 | 9159 | Major tail Protein | Major tail protein [*Klebsiella* phage TSK1] | 212/218 (97) | AXN53791 |
| 8 | 9253 | 9684 | Minor tail protein | Minor tail protein [*Klebsiella* phage Skenny] | 135/143 (94) | QEG07219 |
| 9 | 9674 | 10111 | Neck protein | Putative neck protein [*Klebsiella* phage vB_KpnS_Penguinator] | 144/145 (99) | QEG13398 |
| 10 | 10104 | 10481 | Head-to-tail connector protein | Head-to-tail connector complex protein [*Klebsiella* phage Sweeny] | 119/125 (95) | QEG07140 |
| 11 | 10487 | 10903 | Head-to-tail connector protein | Head-to-tail connector complex protein [*Klebsiella* phage Sin4] | 121/136 (89) | QEG07061 |
| 12 | 10955 | 11263 | Hypothetical protein | Hypothetical protein DDD_7 [*Klebsiella* phage vB_KpnS_KingDDD] | 102/102 (100) | QEG12374 |
| 13 | 11355 | 12296 | Major capsid protein | Major capsid protein [*Klebsiella* phage Sin4] | 264/317 (83) | QEG07059 |
| 14 | 12446 | 12928 | Capsid decoration protein | Capsid decoration protein [*Klebsiella* phage Shelby] | 103/155 (66) | QEG07292 |
| 15 | 12980 | 14110 | Major capsid protein | Putative major capsid protein [*Klebsiella* phage vB_KpnS_IMGroot] | 370/376 (98) | QEG11985 |

| | | | | | | |
|---|---|---|---|---|---|---|
| 16 | 14107 | 14592 | AP2 domain protein | AP2 domain protein [Escherichia phage vB_EcoS_ESCO41] | 130/160 (81) | AQY55280 |
| 17 | 14592 | 15356 | Head morphogenesis protein | Putative head morphogenesis protein [*Klebsiella* phage vB_KpnS_SegesCirculi] | 236/251 (94) | QEG12536 |
| 18 | 15346 | 16665 | Portal protein | Putative portal protein [*Klebsiella* phage vB_KpnS_KingDDD] | 435/439 (99) | QEG12369 |
| 19 | 16712 | 18313 | Terminase large subunit | Terminase large subunit [*Klebsiella* virus GML-KpCol1] | 531/533 (99) | AUE22073 |
| 20 | 18323 | 18847 | Terminase small subunit | Putative terminase small subunit [*Klebsiella* phage GH-K3] | 174/174 (100) | AYP28234 |
| 21 | 18927 | 19190 | Putative structural protein[a] | Hypothetical protein CPT_Shelby_025 [*Klebsiella* phage Shelby] | 85/87 (98) | QEG07286 |
| 22 | 19190 | 19303 | Hypothetical protein | Hypothetical protein [*Klebsiella* phage KpKT21phi1] | 33/36 (92) | AZU98846 |
| 23 | 19521 | 19694 | Hypothetical protein | Hypothetical protein kpv522_21 [*Klebsiella* phage vB_KpnS_KpV522] | 53/56 (95) | AOZ65286 |
| 24 | 19694 | 19927 | Hypothetical protein | Hypothetical protein KLPN1_18 [*Klebsiella* phage KLPN1] | 76/77 (99) | YP_009195358 |
| 25 | 19915 | 20151 | Hypothetical protein | Hypothetical protein [*Klebsiella* virus GML-KpCol1] | 73/78 (94) | AUE22078 |
| 26 | 20221 | 20796 | EaA protein | Putative EaA protein [*Klebsiella* phage GH-K3] | 187/191 (98) | AYP28239 |
| 27 | 20793 | 21005 | Hypothetical protein | Hypothetical protein KPNN141_42 [*Klebsiella* phage KPN N141] | 68/70 (97) | ASW27429 |
| 28 | 21064 | 21276 | Hypothetical protein | Hypothetical protein NJS2_021 [*Klebsiella* phage NJS2] | 59/63 (94) | AXQ67995 |
| 29 | 21285 | 21476 | Hypothetical protein | Hypothetical protein PK_027 [*Klebsiella* phage 1513] | 62/63 (98) | YP_009197834 |
| 30 | 21542 | 21862 | Hypothetical protein | Hypothetical protein KPNN141_40 [*Klebsiella* phage KPN N141] | 105/106 (99) | ASW27427 |
| 31 | 21938 | 22426 | Putative structural protein[a] | Hypothetical protein [*Klebsiella* phage 13] | 151/162 (93) | AZF89865 |

| | | | | | | |
|---|---|---|---|---|---|---|
| 32 | 22426 | 22644 | Hypothetical protein | Hypothetical protein NJS2_016 [*Klebsiella* phage NJS2] | 67/72 (93) | AXQ67990 |
| 33 | 22724 | 22909 | Hypothetical protein | Hypothetical protein KP36_017 [*Klebsiella* phage KP36] | 61/61 (100) | YP_009225979 |
| 34 | 22906 | 23151 | Hypothetical protein | Hypothetical protein GHK3_65 [*Klebsiella* phage GH-K3] | 81/81 (100) | AYP28246 |
| 35 | 23151 | 23453 | Hypothetical protein | Hypothetical protein GHK3_66 [*Klebsiella* phage GH-K3] | 96/100 (96) | AYP28247 |
| 36 | 23450 | 23776 | Hypothetical protein | Hypothetical protein GHK3_67 [*Klebsiella* phage GH-K3] | 104/108 (96) | AYP28248 |
| 37 | 23846 | 24166 | Putative structural protein[a] | Hypothetical protein MezzoGao_55 [*Klebsiella* phage MezzoGao] | 106/106 (100) | ASV45001 |
| 38 | 24177 | 24389 | Hypothetical protein | Hypothetical protein PK_036 [*Klebsiella* phage 1513] | 70/70 (100) | YP_009197843 |
| 39 | 24459 | 24683 | Hypothetical protein | Hypothetical protein [*Klebsiella* phage KP1801] | 73/74 (99) | QHB43310 |
| 40 | 24753 | 24860 | Hypothetical protein | Hypothetical protein NJR15_006 [*Klebsiella* phage NJR15] | 35/35 (100) | AXQ68135 |
| 41 | 24878 | 25057 | Hypothetical protein | Hypothetical protein KLPN1_04 [*Klebsiella* phage KLPN1] | 58/59 (98) | YP_009195344 |
| 42 | 25130 | 25339 | Hypothetical protein | Hypothetical protein MezzoGao_51 [*Klebsiella* phage MezzoGao] | 69/69 (100) | ASV44997 |
| 43 | 25329 | 25481 | Hypothetical protein | Hypothetical protein GHK3_75 [*Klebsiella* phage GH-K3] | 50/50 (100) | AYP28256 |
| 44 | 25554 | 26138 | Putative structural protein[a] | Hypothetical protein [*Klebsiella* phage PhiKpNIH-10] | 188/194 (97) | QHB49641 |
| 45 | 26794 | 27069 | Hypothetical protein | Hypothetical protein GROOT_59 [*Klebsiella* phage vB_KpnS_IMGroot] | 90/91 (99) | QEG12040 |
| 46 | 27072 | 27338 | Hypothetical protein | Hypothetical protein kpv522_78 [*Klebsiella* phage vB_KpnS_KpV522] | 87/88 (99) | AOZ65343 |
| 47 | 27335 | 27664 | Hypothetical protein | Hypothetical protein kpv522_77 [*Klebsiella* phage vB_KpnS_KpV522] | 104/109 (95) | AOZ65342 |

| | | | | | | |
|---|---|---|---|---|---|---|
| 48 | 27747 | 28010 | Hypothetical protein | Hypothetical protein PENG_57 [*Klebsiella* phage vB_KpnS_Penguinator] | 87/87 (100) | QEG13444 |
| 49 | 28083 | 28277 | Hypothetical protein | Hypothetical protein GROOT_55 [*Klebsiella* phage vB_KpnS_IMGroot] | 64/64 (100) | QEG12036 |
| 50 | 28274 | 28513 | Hypothetical protein | Hypothetical protein GROOT_54 [*Klebsiella* phage vB_KpnS_IMGroot] | 76/79 (96) | QEG12035 |
| 51 | 28514 | 28891 | Putative structural protein[a] | Hypothetical protein KOX1_50 [*Klebsiella* phage KOX1] | 125/125 (100) | ARM70374 |
| 52 | 28963 | 29190 | Putative structural protein[a] | Hypothetical protein GROOT_52 [*Klebsiella* phage vB_KpnS_IMGroot] | 74/75 (99) | QEG12033 |
| 53 | 29190 | 29531 | Hypothetical protein | Hypothetical protein DOMN_49 [*Klebsiella* phage vB_KpnS_Domnhall] | 113/113 (100) | QEG11940 |
| 54 | 29524 | 29763 | cytosine DNA methylase | Putative cytosine DNA methylase [*Klebsiella* phage vB_KpnS_KingDDD] | 79/79 (100) | QEG12416 |
| 55 | 29771 | 30463 | DNA methylase | Putative DNA methylase [*Klebsiella* phage vB_KpnS_Alina] | 230/230 (100) | QEG13002 |
| 56 | 30536 | 30820 | Hypothetical protein | Hypothetical protein KOX1_45 [*Klebsiella* phage KOX1] | 68/69 (99) | ARM70369 |
| 57 | 30918 | 31355 | Hypothetical protein | Hypothetical protein GHK3_11 [*Klebsiella* phage GH-K3] | 145/145 (100) | AYP28192 |
| 58 | 31481 | 33049 | Helicase | Helicase [*Klebsiella* phage MezzoGao] | 519/522 (99) | ASV44984 |
| 59 | 33053 | 33514 | Putative ethanolamine utilization protein | Hypothetical protein CPT_Sushi43 [*Klebsiella* phage Sushi] | 152/153 (99) | YP_009196695 |
| 60 | 33584 | 34009 | U-spannin | U-spannin [*Klebsiella* phage Sushi] | 140/141 (99) | YP_009196694 |
| 61 | 34006 | 34488 | Endolysin | Endolysin [*Klebsiella* phage TAH8] | 157/160 (98) | AXQ67962 |
| 62 | 34490 | 34705 | Holin | Holin [*Klebsiella* virus GML-KpCol1] | 71/71 (100) | AUE22114 |
| 63 | 34832 | 35413 | Nucleoside triphosphate hydrolase | Nucleoside triphosphate hydrolase [*Klebsiella* phage Shelby] | 192/193 (99) | QEG07325 |
| 64 | 35410 | 35898 | Polynucleotide kinase | Polynucleotide kinase [*Klebsiella* phage Sushi] | 161/162 (99) | YP_009196690 |
| 65 | 35936 | 37066 | Phosphoesterase | Putative phosphoesterase | 375/376 (99) | ASW27393 |

| | | | | | | |
|---|---|---|---|---|---|---|
| | | | | [*Klebsiella* phage KPN N141] | | |
| 66 | 37162 | 37413 | Hypothetical protein | Hypothetical protein CPT_Sushi36 [*Klebsiella* phage Sushi] | 82/83 (99) | YP_009196688 |
| 67 | 37410 | 37646 | Hypothetical protein | Hypothetical protein CPT_Sushi35 [*Klebsiella* phage Sushi] | 77/78 (99) | YP_009196687 |
| 68 | 37711 | 37947 | Hypothetical protein | Hypothetical protein PK_061 [*Klebsiella* phage 1513] | 78/78 (100) | YP_009197868 |
| 69 | 37951 | 38682 | DNA adenine methyltransferase | DNA adenine methyltransferase [*Klebsiella* phage 1513] | 242/243 (99) | YP_009197869 |
| 70 | 38685 | 38963 | Putative structural protein[a] | Hypothetical protein GHK3_24 [*Klebsiella* phage GH-K3] | 91/92 (99) | AYP28205 |
| 71 | 39037 | 39444 | Holliday junction resolvase | Putative holliday junction resolvase [*Klebsiella* phage Skenny] | 133/135 (99) | QEG07239 |
| 72 | 39444 | 41477 | DNA helicase | DNA helicase [*Klebsiella* phage vB_KpnS_FZ10] | 673/677 (99) | QCG76425 |
| 73 | 41569 | 41970 | Transcriptional regulator | Putative transcriptional regulator [*Klebsiella* phage KLPN1] | 133/133 (100) | YP_009195388 |
| 74 | 42046 | 43005 | DNA primase | DNA primase [*Klebsiella* phage Sweeny] | 316/319 (99) | QEG07158 |
| 75 | 43501 | 44547 | Exonuclease | Exonuclease [*Klebsiella* phage Sushi] | 343/348 (99) | YP_009196679 |
| 76 | 44607 | 45263 | Recombination protein | Recombination protein [*Klebsiella* phage MezzoGao] | 217/218 (99) | ASV44966 |
| 77 | 45300 | 45761 | Single stranded DNA binding protein | Single stranded DNA binding protein [*Klebsiella* phage 1513] | 149/153 (97) | YP_009197877 |
| 78 | 45855 | 48578 | Depolymerase | Depolymerase [*Klebsiella* phage GH-K3] | 903/907 (99) | AYP28213 |
| 79 | 48753 | 50315 | Tail fibre protein | Tail fibre protein [*Klebsiella* phage KP36] | 1147/1266 (91) | YP_009226010 |

*Note*: [a], proteins that were identified by mass spectrometry analysis of MMBB virions and were predicted to be a virion-associated protein by STEP[3], yet had been previously annotated only as “hypothetical proteins”. [b], Protein sequence identity is expressed as a ratio of identical residues/total residues and, in parentheses, as a % identity.

**Supplementary Table 6. Mass spectrometry and STEP$^3$ analysis of *Klebsiella* phage MMNM.**

| Gene | STEP$^3$ | Description |
|---|---|---|
| MMNM_03 | + | Hypothetical protein |
| MMNM_04 | + | Hypothetical protein |
| MMNM_07 | + | Neck protein |
| MMNM_08 | + | Head closure protein |
| MMNM_09 | + | Tail-completion protein |
| MMNM_10 | + | Hypothetical protein |
| MMNM_11 | + | Hypothetical protein |
| MMNM_17 | + | Hypothetical protein |
| MMNM_18 | + | Tape measure protein |
| MMNM_19 | + | Lytic transglycosylase |
| MMNM_20 | + | Hypothetical protein |
| MMNM_21 | + | Baseplate protein |
| MMNM_22 | + | Phospholipase |
| MMNM_23 | + | Baseplate J-like protein |
| MMNM_24 | + | Tail fibre family protein |
| MMNM_26 | + | Base-plate wedge protein |
| MMNM_27 | + | Putative tail-fibre protein |
| MMNM_47 | - | HAD superfamily hydrolase |
| MMNM_50 | - | Polynucleotide kinase (PNK) |
| MMNM_55 | + | Portal protein |
| MMNM_57 | - | Hypothetical protein |
| MMNM_58 | + | Head morphogenesis protein |
| MMNM_65 | + | Coil containing protein |
| MMNM_66 | + | Hypothetical protein |
| MMNM_67 | + | Major capsid protein |

**Supplementary Table 7. Detailed prediction of STEP[3], other available predictors and the BLAST-based baseline predictor on the *Klebsiella* phage MMNM.**

| Gene | BLAST[a] | iVIREONS[b] | PVPred[c] | PVP-SVM[d] | Pred-BVP-Unb[e,*] | PVPred-SCM[f] | STEP[3 g] |
|---|---|---|---|---|---|---|---|
| MMNM_01 | - | 0.709 | 0.045 | 0.092 | 0.005 | 450.560 | 0.072 |
| MMNM_02 | - | -0.418 | 0.085 | 0.070 | 0.013 | 423.970 | 0.218 |
| MMNM_03 | - | -0.717 | 0.915 | 0.600 | 0.890 | 456.590 | 0.750 |
| MMNM_04 | - | 0.878 | 0.898 | 0.878 | 0.130 | 537.390 | 0.932 |
| MMNM_05 | - | -0.961 | 0.010 | 0.005 | 0.024 | 454.120 | 0.116 |
| MMNM_06 | - | -0.942 | 0.000 | 0.004 | 0.004 | 441.330 | 0.057 |
| MMNM_07 | - | -0.497 | 0.573 | 0.684 | 0.404 | 472.100 | 0.689 |
| MMNM_08 | - | 0.916 | 0.311 | 0.443 | 0.159 | 457.410 | 0.736 |
| MMNM_09 | - | 0.781 | 0.695 | 0.418 | 0.645 | 457.670 | 0.706 |
| MMNM_10 | - | 0.938 | 0.476 | 0.695 | 0.967 | 472.660 | 0.968 |
| MMNM_11 | - | 0.995 | 0.563 | 0.711 | 0.918 | 495.130 | 0.873 |
| MMNM_12 | - | 0.832 | 0.144 | 0.095 | 0.002 | 420.030 | 0.088 |
| MMNM_13 | - | -0.783 | 0.004 | 0.013 | 0.002 | 422.750 | 0.106 |
| MMNM_14 | - | -0.997 | 0.337 | 0.394 | 0.105 | 466.370 | 0.352 |
| MMNM_15 | - | 0.573 | 0.215 | 0.278 | 0.018 | 429.450 | 0.083 |
| MMNM_16 | - | -0.645 | 0.630 | 0.516 | 0.261 | 468.390 | 0.466 |
| MMNM_17 | - | -0.427 | 0.926 | 0.840 | 0.564 | 473.380 | 0.849 |
| MMNM_18 | - | 0.976 | 0.749 | 0.737 | 0.562 | 480.270 | 0.907 |
| MMNM_19 | √ | 0.988 | 0.345 | 0.475 | 0.968 | 473.730 | 0.868 |
| MMNM_20 | - | 0.958 | 0.921 | 0.740 | 0.250 | 485.060 | 0.894 |
| MMNM_21 | - | 0.948 | 0.446 | 0.331 | 0.988 | 464.490 | 0.931 |
| MMNM_22 | - | 0.986 | 0.297 | 0.145 | 0.772 | 438.870 | 0.679 |
| MMNM_23 | √ | 0.996 | 0.838 | 0.556 | 0.997 | 477.930 | 0.952 |
| MMNM_24 | √ | 0.975 | 0.432 | 0.461 | 0.961 | 496.040 | 0.925 |

| | | | | | | | |
|---|---|---|---|---|---|---|---|
| MMNM_25 | - | -0.350 | 0.046 | 0.261 | 0.076 | 435.180 | 0.468 |
| MMNM_26 | - | 0.996 | 0.741 | 0.888 | 0.803 | 466.610 | 0.802 |
| MMNM_27 | - | 0.870 | 0.989 | 0.570 | 0.091 | 510.040 | 0.645 |
| MMNM_28 | - | -0.778 | 0.008 | 0.035 | 0.011 | 424.010 | 0.111 |
| MMNM_29 | - | -0.870 | 0.077 | 0.341 | 0.125 | 447.120 | 0.215 |
| MMNM_30 | - | 0.904 | 0.087 | 0.184 | 0.015 | 417.020 | 0.412 |
| MMNM_31 | - | -0.134 | 0.068 | 0.145 | 0.148 | 481.890 | 0.385 |
| MMNM_32 | - | -0.995 | 0.111 | 0.343 | 0.103 | 488.980 | 0.399 |
| MMNM_33 | - | -0.353 | 0.486 | 0.215 | 0.058 | 420.260 | 0.205 |
| MMNM_34 | - | -0.972 | 0.202 | 0.210 | 0.002 | 454.030 | 0.226 |
| MMNM_35 | - | 0.102 | 0.052 | 0.124 | 0.015 | 437.820 | 0.237 |
| MMNM_36 | - | -0.278 | 0.022 | 0.000 | 0.004 | 340.140 | 0.243 |
| MMNM_37 | - | -0.658 | 0.269 | 0.555 | 0.551 | 462.780 | 0.630 |
| MMNM_38 | - | -0.345 | 0.034 | 0.064 | 0.109 | 436.530 | 0.231 |
| MMNM_39 | - | -0.075 | 0.121 | 0.523 | 0.963 | 460.690 | 0.342 |
| MMNM_40 | - | 0.662 | 0.157 | 0.278 | 0.027 | 509.800 | 0.238 |
| MMNM_41 | - | -0.988 | 0.149 | 0.010 | 0.208 | 367.150 | 0.136 |
| MMNM_42 | - | -0.443 | 0.087 | 0.136 | 0.128 | 447.920 | 0.269 |
| MMNM_43 | - | 0.629 | 0.230 | 0.158 | 0.025 | 450.150 | 0.397 |
| MMNM_44 | - | -0.952 | 0.262 | 0.124 | 0.007 | 428.910 | 0.338 |
| MMNM_45 | - | 0.456 | 0.063 | 0.066 | 0.003 | 466.480 | 0.142 |
| MMNM_46 | - | -0.355 | 0.191 | 0.194 | 0.609 | 469.240 | 0.157 |
| MMNM_47 | - | -0.769 | 0.021 | 0.032 | 0.024 | 443.840 | 0.136 |
| MMNM_48 | - | -0.959 | 0.428 | 0.117 | 0.011 | 495.500 | 0.280 |
| MMNM_49 | - | -0.924 | 0.111 | 0.070 | 0.630 | 453.810 | 0.199 |
| MMNM_50 | - | 0.031 | 0.063 | 0.138 | 0.006 | 464.010 | 0.290 |
| MMNM_51 | - | -0.988 | 0.005 | 0.617 | 0.023 | 411.650 | 0.159 |

| | | | | | | | |
|---|---|---|---|---|---|---|---|
| MMNM_52 | - | 0.523 | 0.007 | 0.019 | 0.004 | 423.960 | 0.101 |
| MMNM_53 | - | 0.794 | 0.076 | 0.263 | 0.056 | 439.390 | 0.305 |
| MMNM_54 | - | -0.966 | 0.039 | 0.152 | 0.023 | 428.100 | 0.092 |
| MMNM_55 | - | 0.874 | 0.165 | 0.459 | 0.284 | 455.270 | 0.929 |
| MMNM_56 | - | 0.533 | 0.104 | 0.021 | 0.010 | 432.160 | 0.156 |
| MMNM_57 | - | -0.991 | 0.083 | 0.043 | 0.015 | 400.700 | 0.190 |
| MMNM_58 | - | -0.189 | 0.561 | 0.362 | 0.219 | 454.010 | 0.823 |
| MMNM_59 | - | -0.055 | 0.020 | 0.441 | 0.026 | 443.610 | 0.604 |
| MMNM_60 | - | -0.165 | 0.086 | 0.038 | 0.024 | 417.150 | 0.197 |
| MMNM_61 | - | 0.770 | 0.244 | 0.207 | 0.013 | 468.320 | 0.265 |
| MMNM_62 | - | 0.996 | 0.963 | 0.032 | 0.777 | 501.030 | 0.525 |
| MMNM_63 | - | 0.919 | 0.392 | 0.279 | 0.142 | 453.180 | 0.600 |
| MMNM_64 | - | -0.611 | 0.448 | 0.018 | 0.001 | 454.960 | 0.208 |
| MMNM_65 | - | 0.609 | 0.382 | 0.579 | 0.120 | 457.060 | 0.529 |
| MMNM_66 | - | 0.199 | 0.917 | 0.935 | 0.856 | 502.110 | 0.948 |
| MMNM_67 | - | 0.883 | 0.684 | 0.466 | 0.924 | 474.930 | 0.980 |

*Note*: Proteins identified by mass spectrometry analysis of purified virions are marked in blue. Proteins that were predicted as virion associated proteins by each predictor are marked in green. [a]Proteins that were predicted as virion associated proteins by the BLAST-based predicter were marked with √ or otherwise with -. [b]The prediction score of iVIREONS ranges from -1 to 1. A sequence with predicted score more than 0 is suggested as a virion (structual) protein, according to the description on the iVIREONS server. [c]The prediction cut-off threshold of PVPred is 0.5. A sequence with a prediction score no less than 0.5 is suggested as a virion (structual) protein, which was inferred based on its annotated results. [d]The prediction cut-off threshold of PVP-SVM is 0.45, which was inferred based on its annotated results. [e]The prediction cut-off threshold of Pred-BVP-Unb was set to 0.5. [f]The prediction cut-off threshold of PVPred-SCM is 461.75, according to the description on the PVPred-SCM server. [g]The prediction cut-off threshold of STEP[3] was set to 0.5. *Denotes that Pred-BVP-Unb is not publicly available so was recreated according to published methodology from Arif *et al.*[2].

**Supplementary Table 8. Detailed prediction of STEP[3], other available predictors and the BLAST-based baseline predictor on the *Klebsiella* phage MMBB.**

| Gene | BLAST[a] | iVIREONS[b] | PVPred[c] | PVP-SVM[d] | Pred-BVP-Unb[e,*] | PVPred-SCM[f] | STEP[3 g] |
|---|---|---|---|---|---|---|---|
| MMBB_01 | - | 0.772 | 0.635 | 0.419 | 0.078 | 476.910 | 0.546 |
| MMBB_02 | - | -0.842 | 0.028 | 0.066 | 0.003 | 455.890 | 0.217 |
| MMBB_03 | √ | 0.926 | 0.370 | 0.198 | 0.278 | 455.820 | 0.793 |
| MMBB_04 | √ | 0.874 | 0.697 | 0.922 | 0.807 | 461.750 | 0.783 |
| MMBB_05 | √ | 0.997 | 0.476 | 0.400 | 0.954 | 465.510 | 0.905 |
| MMBB_06 | √ | -0.845 | 0.167 | 0.367 | 0.021 | 430.440 | 0.521 |
| MMBB_07 | - | 0.718 | 0.500 | 0.856 | 0.995 | 472.240 | 0.953 |
| MMBB_08 | - | -0.803 | 0.318 | 0.187 | 0.074 | 429.970 | 0.846 |
| MMBB_09 | - | 0.852 | 0.747 | 0.545 | 0.773 | 469.600 | 0.897 |
| MMBB_10 | - | -0.327 | 0.691 | 0.868 | 0.550 | 486.520 | 0.846 |
| MMBB_11 | - | -0.425 | 0.190 | 0.209 | 0.077 | 444.040 | 0.668 |
| MMBB_12 | - | -0.170 | 0.035 | 0.050 | 0.018 | 392.970 | 0.129 |
| MMBB_13 | - | 0.265 | 0.257 | 0.563 | 0.885 | 443.440 | 0.975 |
| MMBB_14 | - | 0.963 | 0.272 | 0.429 | 0.793 | 488.430 | 0.931 |
| MMBB_15 | - | 0.124 | 0.223 | 0.275 | 0.220 | 448.710 | 0.561 |
| MMBB_16 | - | -0.710 | 0.010 | 0.028 | 0.000 | 427.360 | 0.145 |
| MMBB_17 | - | -0.414 | 0.220 | 0.091 | 0.013 | 462.160 | 0.698 |
| MMBB_18 | - | 0.231 | 0.101 | 0.366 | 0.938 | 443.030 | 0.951 |
| MMBB_19 | - | -0.652 | 0.087 | 0.218 | 0.007 | 428.060 | 0.110 |
| MMBB_20 | - | 0.501 | 0.243 | 0.190 | 0.001 | 449.720 | 0.304 |
| MMBB_21 | - | -0.997 | 0.249 | 0.467 | 0.025 | 430.100 | 0.082 |
| MMBB_22 | - | -0.809 | 0.104 | 0.112 | 0.024 | 454.640 | 0.434 |
| MMBB_23 | - | -0.909 | 0.054 | 0.046 | 0.033 | 413.020 | 0.335 |
| MMBB_24 | - | -0.767 | 0.173 | 0.289 | 0.036 | 413.820 | 0.259 |

| | | | | | | | |
|---|---|---|---|---|---|---|---|
| MMBB_25 | - | 0.816 | 0.253 | 0.350 | 0.058 | 454.970 | 0.411 |
| MMBB_26 | - | -0.872 | 0.006 | 0.100 | 0.001 | 412.840 | 0.110 |
| MMBB_27 | - | -0.973 | 0.023 | 0.007 | 0.029 | 396.230 | 0.208 |
| MMBB_28 | - | -0.629 | 0.067 | 0.074 | 0.165 | 441.720 | 0.245 |
| MMBB_29 | - | -0.954 | 0.859 | 0.957 | 0.180 | 506.680 | 0.422 |
| MMBB_30 | - | -0.955 | 0.029 | 0.252 | 0.103 | 425.270 | 0.217 |
| MMBB_31 | - | -0.991 | 0.094 | 0.211 | 0.023 | 451.290 | 0.327 |
| MMBB_32 | - | -0.984 | 0.008 | 0.103 | 0.018 | 391.850 | 0.137 |
| MMBB_33 | - | -0.998 | 0.442 | 0.022 | 0.417 | 431.780 | 0.290 |
| MMBB_34 | - | -0.343 | 0.322 | 0.542 | 0.026 | 439.710 | 0.243 |
| MMBB_35 | - | -0.235 | 0.029 | 0.048 | 0.014 | 434.090 | 0.201 |
| MMBB_36 | - | -0.196 | 0.251 | 0.140 | 0.031 | 465.340 | 0.439 |
| MMBB_37 | - | 0.412 | 0.198 | 0.246 | 0.075 | 468.890 | 0.868 |
| MMBB_38 | - | -0.845 | 0.048 | 0.078 | 0.022 | 444.320 | 0.226 |
| MMBB_39 | - | -0.149 | 0.251 | 0.013 | 0.027 | 461.530 | 0.186 |
| MMBB_40 | - | -0.584 | 0.006 | 0.005 | 0.004 | 417.030 | 0.109 |
| MMBB_41 | - | -0.998 | 0.546 | 0.021 | 0.012 | 415.410 | 0.149 |
| MMBB_42 | - | -0.647 | 0.021 | 0.053 | 0.138 | 434.370 | 0.507 |
| MMBB_43 | - | -0.566 | 0.490 | 0.380 | 0.022 | 459.290 | 0.396 |
| MMBB_44 | - | -0.468 | 0.294 | 0.087 | 0.058 | 441.050 | 0.536 |
| MMBB_45 | - | -0.304 | 0.081 | 0.093 | 0.015 | 460.930 | 0.165 |
| MMBB_46 | - | -0.986 | 0.000 | 0.029 | 0.007 | 447.110 | 0.190 |
| MMBB_47 | - | -0.525 | 0.026 | 0.113 | 0.575 | 452.330 | 0.149 |
| MMBB_48 | - | -0.590 | 0.012 | 0.064 | 0.003 | 404.360 | 0.122 |
| MMBB_49 | - | -0.078 | 0.836 | 0.528 | 0.027 | 464.190 | 0.731 |
| MMBB_50 | - | -0.394 | 0.198 | 0.005 | 0.006 | 448.760 | 0.202 |
| MMBB_51 | - | 0.981 | 0.883 | 0.720 | 0.050 | 488.660 | 0.550 |

| | | | | | | | |
|---|---|---|---|---|---|---|---|
| MMBB_52 | - | 0.982 | 0.826 | 0.113 | 0.244 | 486.000 | 0.515 |
| MMBB_53 | - | -0.361 | 0.238 | 0.005 | 0.018 | 432.360 | 0.062 |
| MMBB_54 | - | -0.821 | 0.097 | 0.085 | 0.009 | 445.550 | 0.294 |
| MMBB_55 | - | -0.960 | 0.052 | 0.049 | 0.005 | 453.530 | 0.151 |
| MMBB_56 | - | -0.256 | 0.008 | 0.008 | 0.006 | 452.340 | 0.142 |
| MMBB_57 | - | -0.547 | 0.576 | 0.187 | 0.024 | 462.900 | 0.222 |
| MMBB_58 | - | -0.170 | 0.262 | 0.211 | 0.135 | 450.870 | 0.097 |
| MMBB_59 | - | 0.093 | 0.080 | 0.020 | 0.009 | 428.560 | 0.056 |
| MMBB_60 | - | 0.924 | 0.517 | 0.817 | 0.138 | 505.490 | 0.674 |
| MMBB_61 | - | -0.602 | 0.123 | 0.234 | 0.011 | 463.530 | 0.310 |
| MMBB_62 | - | 0.179 | 0.955 | 0.015 | 0.039 | 487.090 | 0.677 |
| MMBB_63 | - | -0.998 | 0.031 | 0.082 | 0.105 | 414.570 | 0.359 |
| MMBB_64 | - | 0.585 | 0.032 | 0.057 | 0.005 | 443.890 | 0.252 |
| MMBB_65 | - | -0.987 | 0.035 | 0.161 | 0.613 | 438.860 | 0.168 |
| MMBB_66 | - | -1.000 | 0.005 | 0.060 | 0.003 | 436.060 | 0.160 |
| MMBB_67 | - | -0.974 | 0.026 | 0.358 | 0.024 | 481.390 | 0.525 |
| MMBB_68 | - | -0.999 | 0.199 | 0.015 | 0.025 | 422.180 | 0.126 |
| MMBB_69 | - | -0.385 | 0.090 | 0.238 | 0.001 | 449.050 | 0.080 |
| MMBB_70 | - | 0.173 | 0.019 | 0.119 | 0.028 | 444.000 | 0.352 |
| MMBB_71 | - | -0.999 | 0.149 | 0.105 | 0.015 | 452.960 | 0.301 |
| MMBB_72 | - | -0.411 | 0.140 | 0.113 | 0.106 | 439.260 | 0.272 |
| MMBB_73 | - | -0.109 | 0.072 | 0.026 | 0.016 | 446.000 | 0.177 |
| MMBB_74 | - | -0.852 | 0.015 | 0.136 | 0.110 | 440.660 | 0.250 |
| MMBB_75 | - | -0.069 | 0.347 | 0.343 | 0.503 | 447.320 | 0.188 |
| MMBB_76 | - | 0.374 | 0.099 | 0.103 | 0.428 | 429.680 | 0.406 |
| MMBB_77 | - | 0.992 | 0.225 | 0.110 | 0.007 | 433.260 | 0.415 |
| MMBB_78 | √ | 0.932 | 0.612 | 0.453 | 0.981 | 482.060 | 0.968 |

| | | | | | | | |
|---|---|---|---|---|---|---|---|
| MMBB_79 | √ | 0.986 | 0.756 | 0.684 | 0.975 | 475.560 | 0.932 |

*Note*: Proteins identified by mass spectrometry analysis of purified virions are marked in blue. Proteins that were predicted as virion associated proteins by each predictor are marked in green. [a]Proteins that were predicted as virion-associated proteins by the BLAST-based predicter were marked with √ or otherwise with -. [b]The prediction score of iVIREONS ranges from -1 to 1. A sequence with predicted score more than 0 is suggested as a virion (structural) protein, according to the description on the iVIREONS server. [c]The prediction cut-off threshold of PVPred is 0.5. A sequence with a prediction score no less than 0.5 is suggested as a virion (structural) protein, which was inferred based on its annotated results. [d]The prediction cut-off threshold of PVP-SVM is 0.45, which was inferred based on its annotated results. [e]The prediction cut-off threshold of Pred-BVP-Unb was set to 0.5. [f]The prediction cut-off threshold of PVPred-SCM is 461.75, according to the description on the PVPred-SCM server. [g]The prediction cut-off threshold of STEP[3] was set to 0.5. *Denotes that Pred-BVP-Unb is not publicly available so was recreated according to published methodology from Arif *et al*.[2].

**Supplementary Table 9. Mass spectrometry and STEP[3] analysis of *Klebsiella* phage MMBB.**

| Gene | STEP[3] | Description |
|---|---|---|
| MMBB_01 | + | Putative tail assembly protein |
| MMBB_03 | + | Putative minor tail protein |
| MMBB_04 | + | Minor tail protein |
| MMBB_05 | + | Tail length tape-measure protein |
| MMBB_06 | + | Tail assembly chaperone |
| MMBB_07 | + | Major tail protein |
| MMBB_08 | + | Minor tail protein |
| MMBB_09 | + | Putative neck protein |
| MMBB_10 | + | Head-to-tail connector protein |
| MMBB_11 | + | Head-to-tail connector complex protein |
| MMBB_13 | + | Major capsid protein/DUF2184 protein |
| MMBB_14 | + | Capsid decoration protein |
| MMBB_15 | + | Major capsid protein |
| MMBB_16 | - | AP2 domain protein |
| MMBB_17 | + | Head morphogenesis protein |
| MMBB_18 | + | Putative portal protein |
| MMBB_21 | - | Hypothetical protein |
| MMBB_31 | - | Hypothetical protein |
| MMBB_52 | + | Hypothetical protein |
| MMBB_60 | + | Membrane protein |
| MMBB_61 | - | Endolysin |
| MMBB_62 | + | Putative holin |
| MMBB_63 | - | Nucleoside triphosphate hydrolase |
| MMBB_64 | - | Polynucleotide kinase (PNK) |
| MMBB_65 | - | Putative phosphoesterase |
| MMBB_70 | - | Hypothetical protein |
| MMBB_76 | - | Putative recombination protein |
| MMBB_78 | + | Depolymerase |
| MMBB_79 | + | Tail fibre protein |

**Supplementary Table 10. Strains, plasmids and primers used in this study.**

| **Strain** | **Description** | **Reference or Source** |
|---|---|---|
| *K. pneumoniae* B5055 | Mouse lethal clinical isolate, K2:O1 | Prof. Richard Strugnell, The University of Melbourne |
| *K. pneumoniae* B5055 *ΔompK36* | B5055 *ΔompK36* | This study |
| *K. pneumoniae* B5055 *Δwzb Δwzc* | B5055 containing a replacement of the *wzb*-*wzc* genes with a kanamycin resistance cassette *ΔwzbΔwzc::*$Km^r$ (non-mucoid) | 3 |
| *K. pneumoniae* ATCC 43816 | Mouse-virulent sequence type 493, K2:O1 | American Type Culture Collection; 4 |
| *K. pneumoniae* AJ094 | Human urinary tract infection isolate, K2 | 5 |
| *K. pneumoniae* AJ097 | Human urinary tract infection isolate, K2 | 5 |
| *K. pneumoniae* AJ210 | Human urinary tract infection isolate, K2/69 | 5 |
| *K. pneumoniae* AJ278 | Human urinary tract infection isolate, K2/13 | 5 |
| *K. pneumoniae* AJ218 | Human urinary tract infection isolate, K54 | 5 |
| *K. pneumoniae* AJ218 *Δwzc* | AJ218 with a mini-Tn5*Km2* transposon insertion within the *wzc* gene (non-mucoid) | 6 |
| **Plasmid** | **Properties** | **Reference** |
| pKD4 | Carries a kanamycin resistance cassette with flanking fragment length polymorphism (FLP) recombinase target (FRT) sites | 7 |
| pACBSR | $Cm^R$, carries L-arabinose inducible I-*Sce*I endonuclease and lambda Red recombination genes | 8 |
| pFLP-BSR | $Cm^R$, carries fragment length polymorphism (FLP) recombinase | 9 |
| **Primers** | **Sequence 5'-3'** | **Reference** |
| ompK36-upF | ctggcagtataaaggctaatggc | 10 |
| ompK36-downR | tgccgctctgattaataacctg | 10 |
| ompK36_pKD4_F | taccggcgttgcgggtgaagctgttgtcgtccagcaggttgattttgtagt gtgtaggctggagctgcttc | This Study |
| ompK36_pKD4_R | taatcagtaagcagtggcataataaaaggcatataacaaacagagggtt acatatgaatatcctccttag | This Study |

**Supplementary Table 11. Genomes of phages used for tree analysis.**

| Phage name | Accession no. |
|---|---|
| **MMNM-related phages in Figure 2a:** | |
| vB KpnM KpV79 | NC_042041.1 |
| vB KpnM FZ14 | MK521906.1 |
| vB KpnM KpV52 | NC_041900.1 |
| 1611E-K2-1 | MG197810.1 |
| vB KpnM IME346 | MK685667.1 |
| vB KpnM 15-38 KLPPOU148 | MN689778.1 |
| PEAT2 | NC_044940.1 |
| MMNM | |
| **MMBB-related phages in Supplementary Figure 2a:** | |
| KOX1 | NC_047825.1 |
| vB KpnS Call | MN013079.1 |
| GH K3 | NC_048162.1 |
| NJS2 | NC_048043.1 |
| MezzoGao | NC_047850.1 |
| MMBB | |
